# Thymus Antibody-Secreting Cells Represent a Tissue Specific Population That Possess an Activated Cellular Phenotype

**DOI:** 10.1101/2022.05.30.494041

**Authors:** KimAnh T. Pioli, Kin H. Lau, Peter D. Pioli

## Abstract

Antibody-secreting cells (ASCs) are key contributors to humoral immunity through immunoglobulin production and the potential to be long-lived. ASCs have been recently identified in the thymus (THY) which is of interest as autoreactive THY B cells regulate T cell tolerance. Here, we showed that the young female THY was skewed towards higher production of ASCs relative to males. However, these differences disappeared with age. Intravenous labeling combined with blockade of CD154(CD40L) suggested that THY ASCs were generated *in situ* in a T cell dependent fashion. Single cell RNA-sequencing revealed that THY ASCs possessed a transcriptional signature indicative of interferon signaling and activation. Flow cytometry confirmed that THY ASCs had increased levels of Toll-like receptor 7 as well as the activation markers CD69 and major histocompatibility class II. Overall, THY ASCs are a unique population that require further investigation to fully appreciate their biological significance.

## Introduction

Humoral, or antibody (Ab) mediated, immunity is essential in preventing infection from a wide variety of bacterium and viruses. This is in part due to the many functions of Ab molecules which include neutralization, opsonization and activation of various effector cell types (Forthal, 2014). Based on namesake alone, it should be no surprise that antibody-secreting cells (ASCs) are key contributors to the humoral immune system. Not only do these cells continuously secrete Abs (Slifka et al., 1998) but a segment of the population has been shown to be long-lived in both mice (Chernova et al., 2014; Manz et al., 1998; Manz et al., 1997; Slifka *et al*., 1998) and humans (Amanna et al., 2007; Halliley et al., 2015; Landsverk et al., 2017).

Aside from the ability to produce Abs, ASCs possess numerous immunomodulatory roles which can be at least partially attributed to their ability to secrete cytokines (Fillatreau, 2019; Pioli, 2019; Wang et al., 2021). For instance, IL-10 production by ASCs has been associated with the suppression of neuroinflammation (Rojas et al., 2019) as well as attenuation of the response to *Salmonella* infection (Lino et al., 2018). In the latter case, this phenotype has been linked with surface expression of LAG-3 (Lino et al., 2018). In contrast, ASCs have also demonstrated production of proinflammatory cytokines such as tumor necrosis factor-alpha (TNF-α) (Fritz et al., 2011) and interleukin-17 (IL-17) (Bermejo et al., 2013) which play roles in gut homeostasis and response to *Trypanosoma cruzi* infection, respectively.

The continual exploration of ASC biology has led to the discovery of regulatory roles which reach beyond the immune response. Along these lines, it now appreciated that ASCs influence hematopoiesis with this area being most well studied in the bone marrow (BM). Multiple reports have demonstrated the ability of plasma cells (PCs), which are post-mitotic mature ASCs, to regulate not only the numerical production of myeloid cells but also the type of myeloid progeny being produced (Meng et al., 2019; Pioli et al., 2019). Interestingly enough, the ability to drive myeloid expansion appears to be critically dependent on the “age” of PCs. This was exemplified in co-culture experiments in which BM PCs from old mice (17+ months (mo.)) enhanced myeloid output from co-cultured hematopoietic stem cells (HSCs) (Pioli et al., 2019). This function was lacking from PCs isolated the BM of young (2-3 mo.) mice. In contrast, BM PCs from young mice possessed the potential to promote B lymphopoiesis when cultured alongside common lymphoid progenitors (CLPs) (Pioli et al., 2019). Together, these findings demonstrate a critical role for ASCs in regulating blood cell development. However, whether ASCs regulate hematopoietic processes in other organs remains to be determined.

Recent studies in humans have identified ASC populations in the neonatal THY which are preserved and potentially expanded over the course of aging (Cordero et al., 2021; Nunez et al., 2016). Histological analyses in humans have shown these cells to be located within the THY perivascular space (PVS) (Cordero et al., 2021; Nunez et al., 2016) and recent RNA-sequencing (RNA-seq) experiments have inferred an intra-THY development of these cells (Cordero et al., 2021). This latter point being of particular interest as experiments using mice have shown the ability of autoreactive THY B cells to class-switch following their interactions with developing T lymphocytes (Perera et al., 2016). That being said, a comprehensive analysis of THY ASCs and how they compare to their peripheral counterparts is lacking.

In this report, we showed that the production of THY ASCs favored the female sex in youth (3 mo. old mice) and reached equivalence between the sexes by middle age (12 mo. old). Using intravenous antibody labeling as well as *in vivo* blockade of CD154(CD40L) signaling, we demonstrated that ASCs from both sexes were produced locally in the THY and required T cell-derived instructive signals for their generation. Similar to their peripheral counterparts, THY ASCs also required the expression of BLIMP-1 as its deletion ablated THY ASCs. Using single cell RNA-seq (scRNA-seq), we identified three developmental lineages which were shared between ASCs isolated from the BM, spleen (SPL) and THY. However, closer inspection of VDJ and transcriptional signatures revealed an immunoglobulin repertoire and interferon gene signature that was unique to THY ASCs and was highlighted by increased expression of genes such as *Tlr7* and *Ly6c2*. Overall, these studies provide a valuable resource to be leveraged for the generation of hypotheses and future exploration of THY ASCs.

## Results

### Young female mice have increased numbers of thymus antibody-secreting cells relative to males

Previous work has identified the presence of ASCs in the human THY (Cordero et al., 2021; Nunez et al., 2016). Similarly, ASCs have been observed in the mouse THY (Haba and Nisonoff, 1992; Hidalgo et al., 2020) but remain to be fully characterized. To begin to address this deficiency, we euthanized 3 months (mo.) old Prdm1-enhanced yellow fluorescence protein (eYFP) reporter mice and examined ASCs in the THY. These mice possess a bacterial artificial chromosome (BAC) that expresses enhanced yellow fluorescent protein (eYFP) downstream of the *Prdm1* gene (Fooksman et al., 2014) and provide a suitable means to identify ASCs when combined with the surface expression of selected proteins. We identified ASCs as CD138^HI^ IgD^-/LO^ CD90.2^-/LO^ Prdm1-eYFP^+^. Subsequently, B220 expression (Chernova et al., 2014) further subdivided the ASC population into B220^+^ plasmablasts (PBs) and B220^-^ PCs (**Figure 1A**). PBs and PCs represent relatively short-lived proliferative and post-mitotic mature ASC populations, respectively. In agreement with previous studies (Gui et al., 2011; Min et al., 2006), 3 mo. old female mice had increased overall cellularity in the THY compared to age-matched males (**Figure 1B**). Percentages (**Figure 1C**) and numbers (**Figure 1D**) of both PBs and PCs were both significantly increased in female THY relative to male mice. Furthermore, the female THY was skewed towards a PB phenotype (**Figures 1C-1D**) perhaps indicative of a heightened level of immune reactivity in the female THY. The BM and SPL did not demonstrate similar phenotypes (**Figures S1A-S1F**) lending specificity to our observations in the THY.

**Figure 1.**
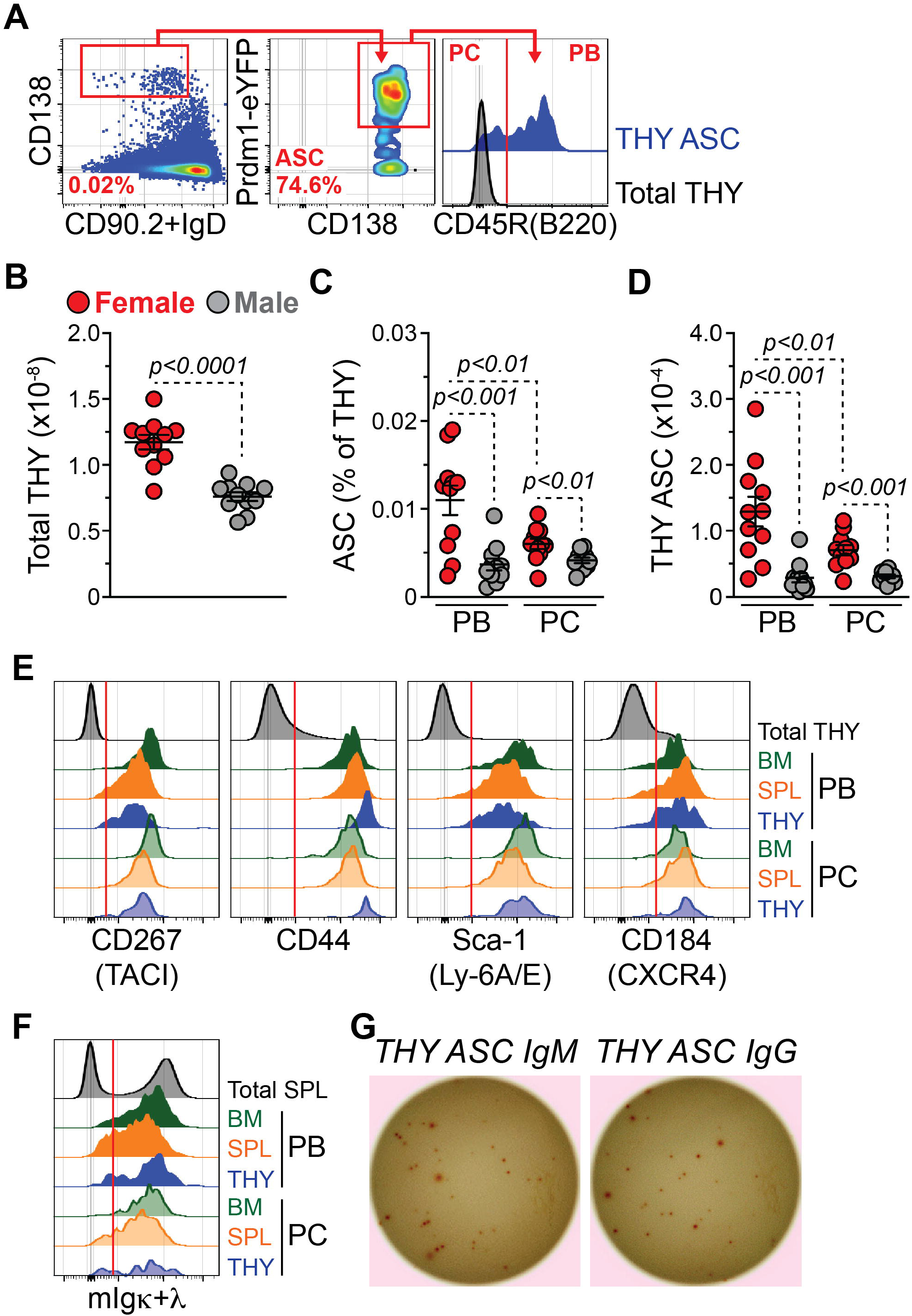
Thymus antibody-secreting cells are preferentially increased in young female thymus. (A) Flow cytometry plots depicting representative gating of total ASCs as well as PBs and PCs from the THY. Cells pre-gated on total singlets. Red vertical line added to CD45R(B220) histogram to show cut-off for positive staining. Numbers in plots represent percentages within the immediate parent population. (B) Total THY cell numbers. (C) Percentages of THY PBs and PCs. (D) Numbers of THY PBs and PCs. (E) Flow cytometry overlays showing CD267(TACI), CD44, Sca-1(Ly-6A/E), and CD184(CXCR4) staining on the surface of PBs and PCs from BM, SPL and THY. Total THY is shown for comparison. Red vertical lines added to histograms to show cut-offs for positive staining. (F) Flow cytometry overlays showing mIgκ+λ staining on the surface of PBs and PCs from BM, SPL and THY. Total SPL is shown for comparison. Red vertical line added to histogram to show cut-off for positive staining. (G) Representative images of IgM and total IgG THY ASC ELISpots. 100 cells per well were plated in triplicate. Data shown for males. Experiments were reproduced with female ASCs. (B-D) Symbols represent individual mice. Female: n = 11; Male: n = 11. Horizontal lines represent mean ± SEM. Unpaired Student’s t-Test for inter-sex comparisons. Paired Student’s t-Test for intra-sex comparisons.

Further characterization of THY ASCs, relative to those in BM and SPL, demonstrated surface expression of ASC-associated proteins such as CD267(TACI) (Cheng et al., 2020; Shen et al., 2014), CD44 (Matsumoto et al., 2014; Suzuki-Yamazaki et al., 2017; Underhill et al., 2002), Sca-1(Ly-6A/E) (Wilmore et al., 2017) and CD184(CXCR4) (Shen et al., 2014) (**Figure 1E**). In addition, THY ASCs expressed membrane immunoglobulin (mIgκ+λ) like that previously observed in ASCs from the BM and SPL (**Figure 1F**) (Blanc et al., 2016; Pinto et al., 2013; Pioli et al., 2019). In some instances, organ-to-organ expression differences were evident (**Figures S1G-S1K**). For example, BM ASCs expressed the highest amounts of CD267(TACI) as assessed by flow cytometric geometric mean fluorescence intensity (gMFI) (**Figure S1G**). In contrast, THY ASCs expressed the highest levels of CD44 (**Figure S1H**). Specific to PCs, the THY had modestly, but significantly, increased levels of CD184(CXCR4) expression (**Figure S1J**).

To functionally validate THY ASCs, we purified them via fluorescence-activated cell sorting (FACS) (**Figure S2**) and performed enzyme-linked immunosorbent spot (ELISpot) assays for IgM and total IgG (**Figure 1G**). As expected, THY ASCs were able to secrete both IgM and total IgG when cultured overnight (**Figure 1G**) confirming them to be *bona fide* ASCs. Overall, these data identified ASCs in the homeostatic mouse THY and demonstrated an organ-specific, sex-biased expansion of ASC populations in the female THY.

### Age-associated accumulation of thymus antibody-secreting cells is sex dependent

Aging in mice has been associated with the accumulation of ASCs in tissues such as BM and SPL (Benet et al., 2021; Lino et al., 2018; Pioli et al., 2019). Furthermore, ELISpot data from humans is indicative of increased ASCs in the THY with age (Nunez et al., 2016). To determine if percentages and/or numbers of mouse THY ASCs fluctuate with age, we performed flow cytometry on THY from middle-aged (12 mo. old) Prdm1-eYFP mice and compared those data to our 3 mo. old data set (**Figures S3A-S3F**). As controls for age-associated ASC accumulation, we also assessed BM and SPL (**Figures S3G-S3P**). As expected, THY cellularity decreased with age (**Figure 3SB**) and this was accompanied by a general increase in the percentages of PBs and PCs in the THY of both sexes (**Figures 3SC-3SD**). However, male THY demonstrated increased numbers of PBs and PCs with age that was not apparent in females (**Figures S3E-S3F**). Notably, percentages and numbers of PBs and PCs increased in the BM and SPL with age for both sexes (**Figures S3H-S3K, S3M-S3P**). In most instances, BM and SPL ASC populations showed the greatest accumulation in females compared to males. In summary, these analyses showed that THY ASCs undergo a distinct pattern of aging when compared to the BM and SPL.

### Thymus antibody-secreting cells are not recently immigrated thymic populations

B cells have been observed in the THY of both humans (Isaacson et al., 1987) and mice (Miyama-Inaba et al., 1988). Using parabiosis experiments, it was suggested that these were mostly non-circulatory and thus were THY-derived (Perera et al., 2013; Perera et al., 2016). This concept is supported by the previous identification of B cell progenitors within the THY (Mori et al., 1997). Collectively, these studies raise the possibility that THY ASCs may be generated locally within the THY rather than a result of immigration from the periphery.

To address this idea, we injected phosphate buffered saline (PBS) or 1 μg of αCD45-PE Abs intravenously (i.v.) into 3 mo. old Prdm1-eYFP mice (**Figure 2A**). After 5 minutes (min.) or 24 hours (hr.), we euthanized mice and analyzed the distribution of αCD45-PE staining within the peripheral blood (BLD), BM, SPL and THY. αCD45-PE labeled cells within a tissue are indicative of those localized within the tissue vasculature or having trafficked within the bloodstream during the labeling period (Anderson et al., 2014; Ruscher and Hogquist, 2018; Thanabalasuriar et al., 2016). As expected, cells from the BLD of mice infused with αCD45-PE for 5 min. possessed >99% labeling demonstrating the efficacy of our protocol (**Figure 2B**).

**Figure 2.**
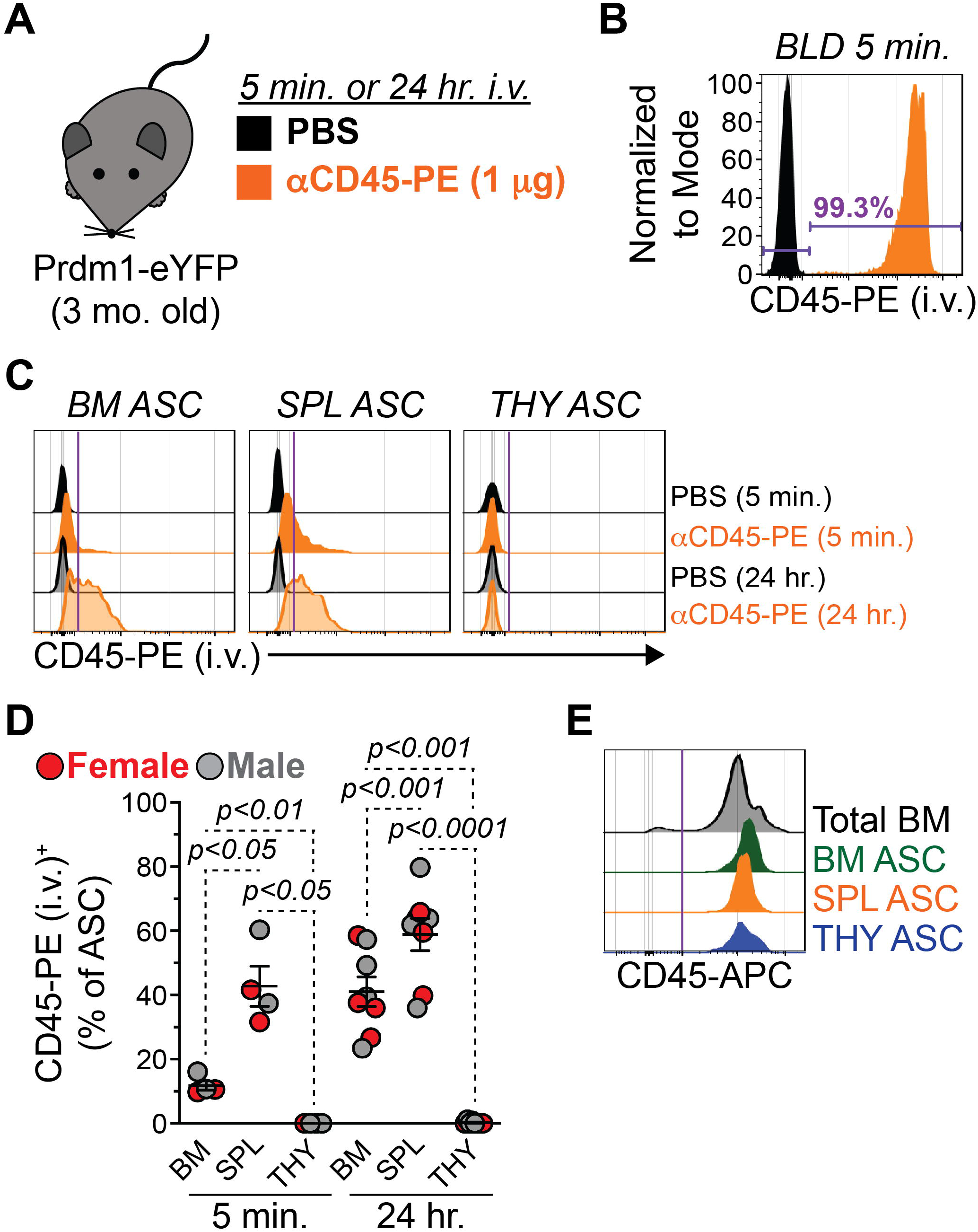
Thymic antibody-secreting cells are not recent thymic immigrants. (A) Young (3 mo. old) female and male Prdm1-eYFP mice received i.v. injections through the retro-orbital (r.o.) sinus containing either PBS or αCD45-PE Abs (1 μg). Animals were euthanized and assessed for αCD45-PE labeling 5 min. or 24 hr. post-injection. (B) Flow cytometry histograms overlaying CD45-PE fluorescence of total BLD cells from animals that received either PBS or αCD45-PE for 5 min. Number in plot indicates percentage of total BLD cells positive for CD45-PE (i.v.) staining 5 min. post-injection of αCD45-PE. (C) Flow cytometry plots depicting αCD45-PE (i.v.) labeling of ASCs from BM, SPL and THY. Staining from PBS-treated mice and mice that received αCD45-PE Abs for 5 min. or 24 hr. is shown for comparison. Purple vertical lines added to CD45-PE (i.v.) histograms to show cut-off for positive staining. (D) Percentages of BM, SPL and THY ASCs labeled with αCD45-PE 5 min. or 24 hr. post-injection. Symbols represent individual mice. 5 min.: n = 4; 24 hr.: n = 8. Horizontal lines represent mean ± SEM. One-way ANOVA with Tukey’s correction. (E) Flow cytometry histogram showing CD45-APC *ex vivo* staining of total BM cells and ASCs from BM, SPL and THY. Purple vertical line added to CD45-APC histogram to show cut-off for positive staining.

αCD45-PE labeling of ASCs revealed organ specific staining patterns (**Figure 2C**). In agreement with previous studies (Chernova et al., 2014), BM ASCs demonstrated low levels of labeling (∼11.8%) after 5 min. while that of SPL ASCs was more extensive (∼42.7%) (**Figure 2D**). In the THY, 0% of ASCs were labeled after 5 min (**Figure 2D**). Using an extended 24 hr. labeling period, we observed high percentages of αCD45-PE^+^ ASCs in the BM (∼41.0%) and SPL (∼58.8%); however, THY ASCs remained unlabeled (**Figure 2D**). The lack of αCD45-PE staining on THY ASCs was not due to absence of CD45 as *ex vivo* staining with αCD45-APC demonstrated similarities in CD45 expression amongst ASCs from all 3 organs analyzed (**Figure 2E**). These experiments suggested that THY ASCs were not the result of recent immigration and were most likely generated from a THY-localized immune response.

### Expression of BLIMP-1 and CD154(CD40L)-derived signals are required for thymus antibody-secreting cell production

It is well known that BLIMP-1, encoded for by *Prdm1*, is a key regulator of ASC differentiation and function (Minnich et al., 2016; Shaffer et al., 2002; Tellier et al., 2016). In fact, the genetic ablation of *Prdm1* in B cells leads to essentially a complete loss of ASCs in the SPL following immunization (Shapiro-Shelef et al., 2003). To determine the importance of BLIMP-1 for the formation of THY ASCs, we bred mice with a conditional allele of *Prdm1* (Shapiro-Shelef et al., 2003) to a strain possessing the B cell specific Mb1(Cd79a)-Cre (Hobeika et al., 2006). For these experiments, we performed analyses using 3-5 mo. old mice which were not age matched. In the absence of the Prdm1-eYFP reporter, THY ASCs were identified as CD138^HI^ IgD^-/LO^ CD90.2^-/LO^ CD267(TACI)^+^ CD44^+^ (**Figure 3A**). As expected, mice with the *Prdm1*^Fl/Fl^ *Mb1*^+/+^ genotype were wildtype (WT) and possessed a distinct ASC population (**Figure 3A**). This population was lost in *Prdm1*^Fl/Fl^ *Mb1*^+/Cre^ ASC knockout (KO) mice (**Figures 3A-3B**) demonstrating the conserved reliance on BLIMP-1 for the production of THY ASCs.

**Figure 3.**
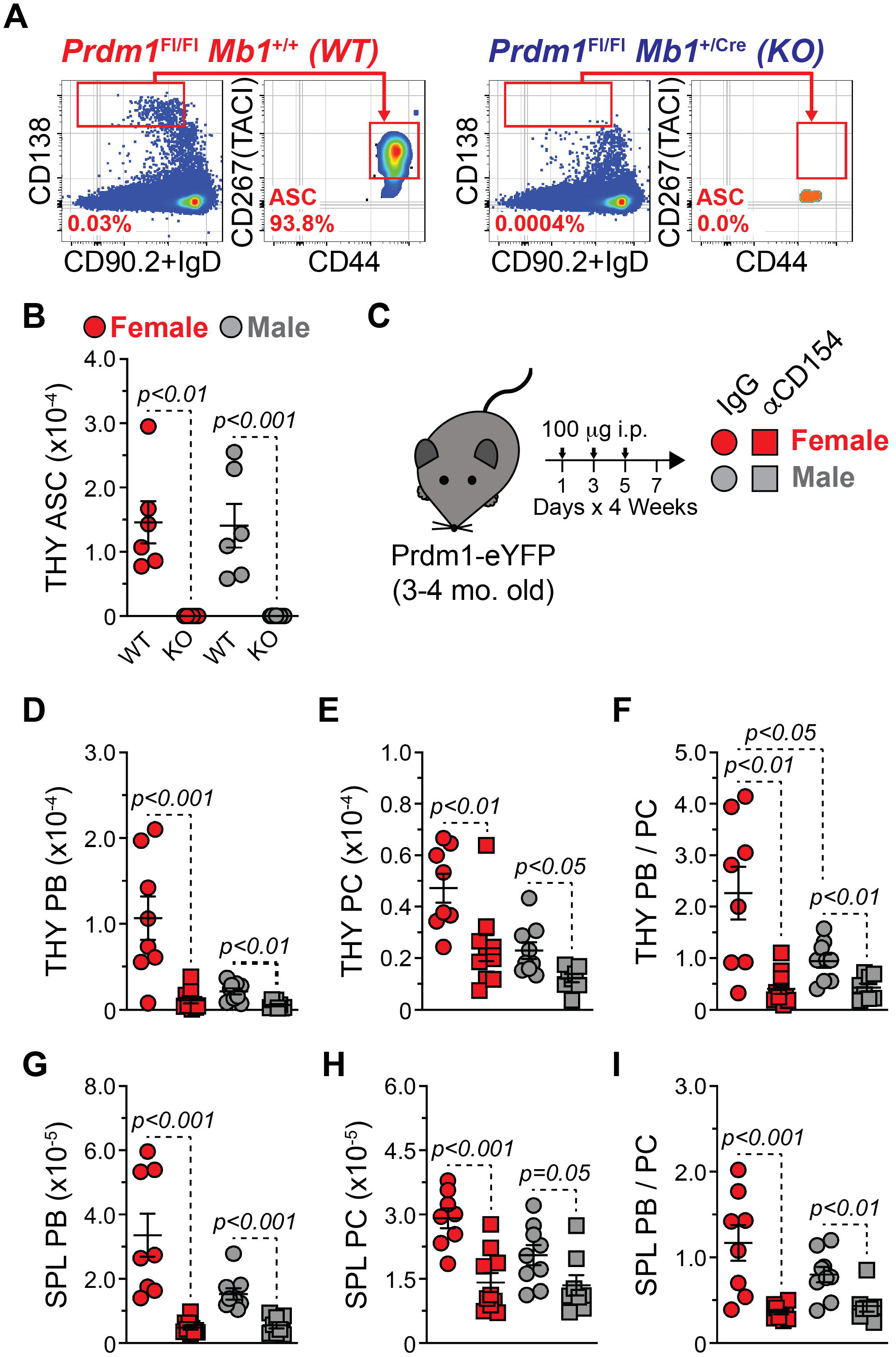
Expression of BLIMP-1 and CD154(CD40L)-derived signals are required for thymus antibody-secreting cell production. (A) Flow cytometry plots showing representative gating of THY ASCs from *Prdm1*^Fl/Fl^ *Mb1*^+/+^ WT and *Prdm1*^Fl/Fl^ *Mb1*^+/Cre^ KO mice. Numbers in plots represent percentages within the immediate parent population. (B) Numbers of THY ASCs from 3-5 mo. old Prdm1 WT and KO mice. (C) Young (3-4 mo. old) Prdm1-eYFP female and male mice received 1200 μg total of hamster IgG isotype control or anti-mouse CD154(CD40L) (MR-1) Abs. Administration was split evenly amongst 12-total i.p. injections (100 μg per injection, 100 μl volume) over a 4-week span. (D-E) Numbers of THY (D) PBs and (E) PCs in IgG- and αCD154-treated mice. (F) Ratio of THY PB / PC in IgG- and αCD154-treated mice. (G-H) Numbers of SPL (G) PBs and (H) PCs in IgG- and αCD154-treated mice. (I) Ratio of SPL PB / PC in IgG- and αCD154-treated mice. (B, D-I) Symbols represent individual mice. Horizontal lines represent mean ± SEM. Unpaired Student’s t-Test. (B) Female WT: n = 6; Female KO: n = 6; Male WT: n = 6; Male KO: n = 8. (D-I) Female IgG: n = 8; Female αCD154: n = 10; Male IgG: n = 9; Male αCD154: n = 8. Unpaired Student’s t-Test.

Aside from intrinsic regulators (i.e., BLIMP-1), the surrounding milieu in which a B cell is activated can influence ASC differentiation. For example, ASCs can be generated in the context of either a T cell independent or T cell dependent immune response (Nutt et al., 2015). Recently, it has been shown that THY B cells class switch when activated in the presence of T cell-derived signals and genetic ablation of TCRα, MHC II or CD40 all led to a significant loss of class switched THY B cells (Perera et al., 2016). As such, we hypothesized that inhibition of B cell-T cell interactions would lead to a reduction in THY ASCs. To test this, cohorts of 3-4 mo. old female and male Prdm1-eYFP mice were treated intraperitoneally (i.p.) with isotype control IgG or αCD154(CD40L) (MR-1) blocking Abs over the course of 1 month (**Figure 3C**). Subsequently, mice were euthanized and ASC populations were assessed. Upon blockade of CD154(CD40L) signals, we observed a significant reduction in the numbers of THY PBs and PCs (**Figures 3D-3E**). Ratios of PBs to PCs (PB / PC) were also significantly reduced in the THY of both female and male mice that received αCD154(C40L) Abs compared to those administered isotype control IgG (**Figure 3F**). This indicated a greater numerical loss in the PB population. Examination of the SPL (**Figures 3G-3I**) yielded largely similar results supporting a common requirement, at least to some degree, for T cell-derived signals to produce ASCs in both organs.

### In vivo CD40 activation does not augment formation of thymus antibody-secreting cells

Given the requirement for CD154(CD40L), we wanted to know whether ectopic CD40L:CD40 signals would be sufficient to drive expansion of THY ASCs. 3-4 mo. old female and male Prdm1-eYFP mice were treated i.p. with isotype control IgG2a or αCD40 agonistic Abs on days -1 and 0 (**Figure 4A**). Mice were subsequently euthanized on days 2, 4, 6 and 28 post-injection to assess production of PBs and PCs in both the THY (**Figures 4B-4E**) and SPL (**Figures 4F-4I**). For these analyses, isotype control treated mice from different days were pooled due to no clear variation between timepoints. Enumeration of THY PBs (**Figures 4B-4C**) and PCs (**Figures 4D-4E**) showed little fluctuation over the time course.

**Figure 4.**
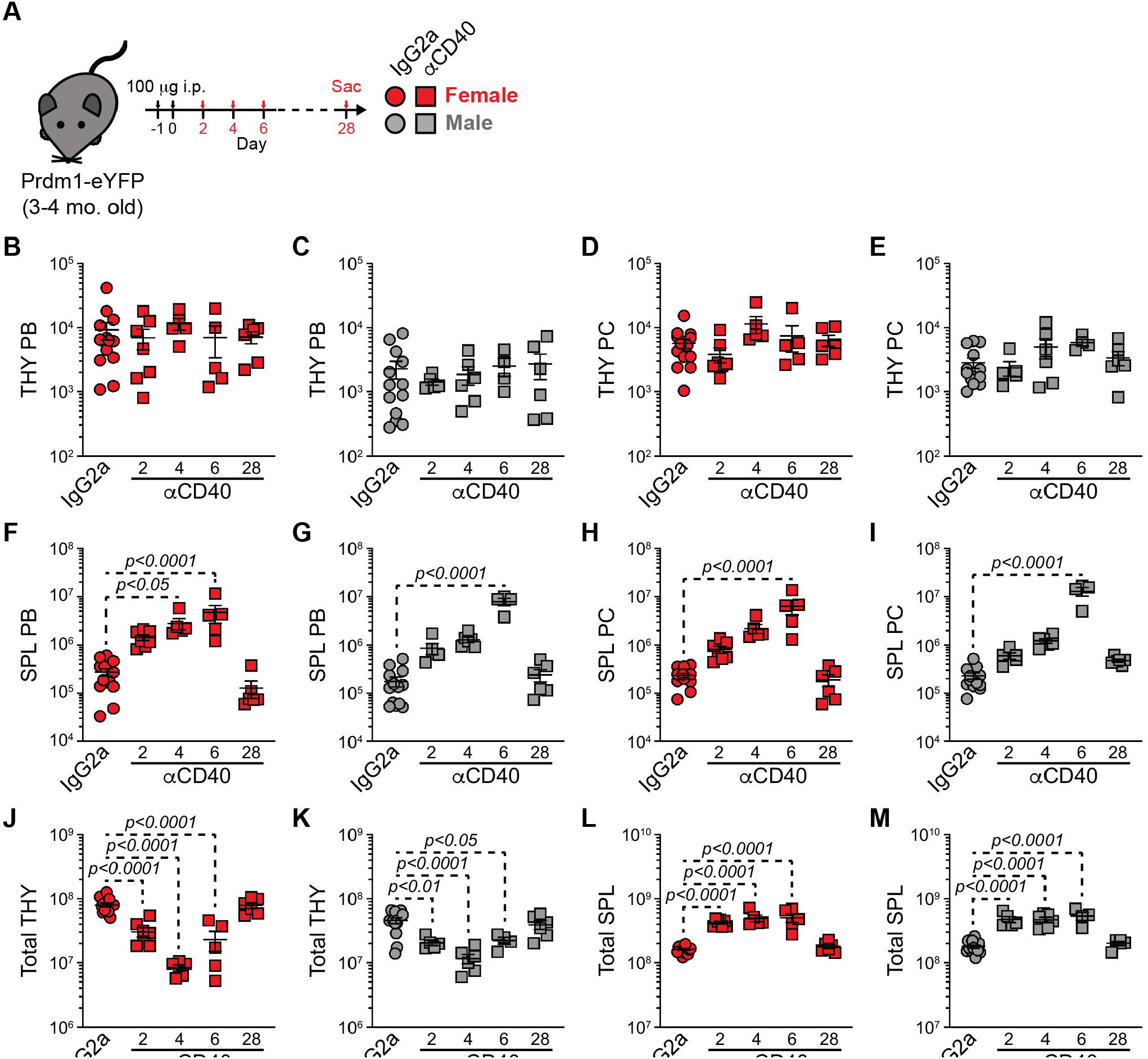
Ectopic CD40 stimulation does not increase thymus antibody-secreting cell numbers. (A) Young (3-4 mo. old) Prdm1-eYFP female and male mice received a total of 200 μg of rat IgG2a isotype control or anti-mouse CD40 (FGK4.5/FGK45) antibodies. Doses (100 μg per injection, 100 μl volume) were administered i.p. on days -1 and 0. Animals were euthanized on days 2, 4, 6 and 28 post-injection and ASCs were analyzed. (B-C) Numbers of THY PBs in (B) female and (C) male IgG2a- and αCD40-treated mice. (D-E) Numbers of THY PCs in (D) female and (E) male IgG2a- and αCD40-treated mice. (F-G) Numbers of SPL PBs in (F) female and (G) male IgG2a- and αCD40-treated mice. (H-I) Numbers of SPL PCs in (H) female and (I) male IgG2a- and αCD40-treated mice. (J-K) Total THY cell numbers in (J) female and (K) male IgG2a- and αCD40-treated mice. (L-M) Total SPL cell numbers in (L) female and (M) male IgG2a- and αCD40-treated mice. (B-M) Symbols represent individual mice. Horizontal lines represent mean ± SEM. IgG2a data represent pooled control mice from days 2, 4, 6 and 28. Female IgG2a: n = 13; Female αCD40 day 2: n = 7; Female αCD40 day 4: n = 5; Female αCD40 day 6: n = 5; Female αCD40 day 28: n = 6; Male IgG2a: n = 13; Male αCD40 day 2: n = 5; Male αCD40 day 4: n = 6; Male αCD40 day 6: n = 4; Male αCD40 day 28: n = 6. One-way ANOVA with Dunnett’s correction. All comparisons made against IgG2a controls.

In contrast, both SPL PBs (**Figures 4F-4G**) and PCs (**Figures 4H-4I**) were increased, in particular at day 6 of the response. It should be noted that no long-term (day 28) alterations of PB and PC numbers were observed. Further examination of total cellularity revealed significant THY atrophy between days 2-6 for both sexes which rebounded by day 28 (**Figures 4J-4K**). This was opposite to the SPL which demonstrated a short-term expansion in cell numbers (**Figures 4L-4M**) and may have contributed to the inability to boost ASC numbers in the THY. Regardless, these data suggested that αCD40 ligation *in vivo* can transiently increase PBs and PCs. However, the ability to do so appeared to be tissue restricted as ectopic CD40 stimulation alone could not expand THY ASCs.

### Antibody-secreting cell subsets exist with divergent gene ontology associations

The bulk of the proceeding data indicated that THY ASCs were potentially divergent when compared to those from the BM and SPL. To better understand these differences, we FACS-purified ASCs (**Figure S2**) from the BM, SPL and THY and performed scRNA-seq to assess single cell gene expression (scGEX) as well as VDJ (scVDJ) usage via the 10x Genomics platform (**Figure 5A**). This was done with cells separately isolated from pools of 3 female and 3 male 3-months old Prdm1-eYFP mice. Similar to what has previously been shown (Scharer et al., 2020), expression of *Xbp1* and *Ell2* was readily detectable amongst ASCs (**Figure 5B**) providing validation of cell identity. Clustering of ASCs from the 6 total samples led to the classification of 12 clusters labeled 0-11 (**Figure 5C**). While there was some variability between the sexes regarding cluster contribution to total ASCs, Cluster 0 was a consistent majority within the THY of both female and male mice (**Figure 5D**). In contrast, BM and SPL showed a greater prevalence for “minor” clusters such as Cluster 4 (**Figure 5D**).

**Figure 5.**
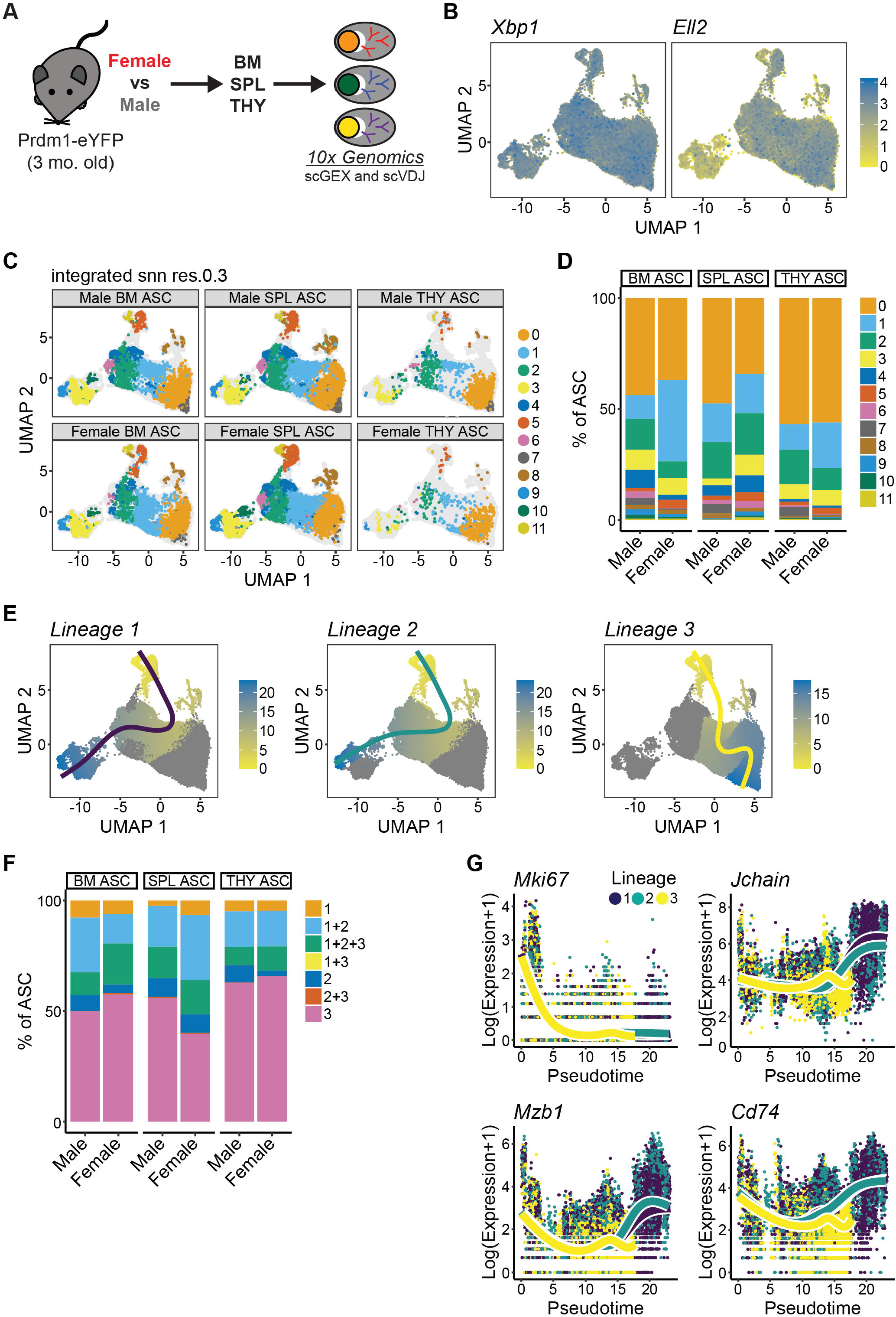
Single cell RNA sequencing identifies antibody-secreting cell clusters and divergent lineages that exist in bone marrow, spleen and thymus. (A) Schematic of scRNA-seq analysis of ASCs from female and male BM, SPL and THY. Subsequent data derived from pools of 3 female and 3 male mice, respectively. (B) UMAP plots showing log-normalized unique molecular identifier (UMI) counts for *Xbp1* and *Ell2*. (C) UMAP plots showing single ASCs and cluster assignments from female and male BM, SPL and THY. (D) Graph depicting relative cluster contribution to ASC pools from female and male BM, SPL and THY. (E) Pseudotime plots showing the identification of 3 ASC lineages. (F) Relative contributions of various pseudotime lineages to male and female ASCs from BM, SPL and THY. (G) Plots illustrating changes in Log(Expression+1) of *Mki67, Jchain, Mzb1* and *Cd74* over the course of pseudotime. Individual symbols depict single cells within a particular lineage.

To better understand transcriptional differences between clusters, we utilized the *scran* package to identify marker genes that were significantly enriched in a cluster as indicated by an area under the curve (AUC) > 0.75 and false discovery rate (FDR) < 0.05 (**Table S1**). Using this method, the AUC represents the probability that expression of a given gene from a cell in a particular cluster is greater than any random cell from another cluster. When comparing clusters, Cluster 11 possessed the most marker genes (1545 total, **Figure S4A, Table S1**). Interestingly, we observed a dichotomy in terms of the contribution of Ig versus non-Ig genes to the total number of markers. For example, Clusters 0-2 and 7 were heavily skewed towards Ig marker genes while in contrast, Clusters 3 and 9-11 showed dominance by non-Ig genes (**Figure S4A**). Previous research has shown that ASC heavy chain isotypes coincide with particular gene programs (Price et al., 2019). Accordingly, Clusters 0, 1, 5 and 7 were significantly enriched for *Ighm, Ighg2b, Ighg2c* and *Igha* isotypes (**Table S1**). Clusters 4, 6 and 8 were correlated with *Ighg2b, Ighg2c* and *Igha* while *Ighg2c* and *Igha* were enriched for in Cluster 2 (**Table S1**). Notably, Clusters 3 and 9-11 did not demonstrate significant association with a particular IgH constant region gene (**Table S1**).

Non-Ig cluster marker genes were assessed for gene ontology associations using the Metascape on-line tool (Zhou et al., 2019) (**Table S2**). For these analyses, we focused on GO Biological Processes and only considered terms which possessed a gene overlap of ≥ 10. Clusters 0-2 and 7 did not demonstrate any term association due to a low number of non-Ig marker genes (**Figure S4A, Table S1**). However, we identified multiple instances where clusters shared a certain degree of similarity. For example, Clusters 3 and 8-11 shared enrichment for protein-related biological processes such as cytoplasmic translation (GO:0002181), translation (GO:0006412) and those related to ribosome production (GO:0022613, GO:0042254, GO:0042255) (**Figure S4B, Table S2**). Metabolic biological processes were enriched for in many clusters (**Figure S4C, Table S2**) and included terms such as oxidative phosphorylation (GO:0006119) and aerobic respiration (GO:0009060) which was in agreement with what is known regarding ASC metabolism (Lam et al., 2016; Lam et al., 2018; Price et al., 2018). However, gene ontology analyses did identify some distinguishing features amongst the clusters with Cluster 11 showing a clear enrichment for RNA processing-related terms (**Figure S4D, Table S2**) and processes related to cell proliferation (mitotic cell cycle process, GO:1903047) (**Figure S4E, Table S2**). In many ways, Cluster 11 in our dataset resembled previous work which identified a “mitotic” ASC cluster within sorted B220^+^ 2-NBDG^-^ SPL ASCs (Lam et al., 2018). In fact, many of the clusters and associated ontologies that we observed shared commonality with earlier single-cell ASC studies (Lam et al., 2018) lending validity to our analyses.

### Antibody-secreting cells are a composite of divergent lineages which display organ-specific changes in gene expression

The above analyses suggested some fundamental differences existed between ASC clusters. In some instances, these differences were related to UMAP position. A prime example of this was Clusters 0-2 and 7 that possessed minimal non-Ig marker genes, lacked significant gene ontology enrichment and were segregated on the right portion of the UMAP plots (**Figure 5C**). This suggested that ASCs might have divergent lineages or “fates”. To investigate this further, we performed pseudotime analysis using the *tradeSeq* package (Van den Berge et al., 2020). Lineages were rooted (pseudotime = 0) at Cluster 11. This cluster was chosen based on its gene ontology association with mitosis (**Figure S4E, Table S2**) as well as high expression of cell cycle genes such as *Ccna2* (**Figure S4F**). Pseudotime analysis revealed 3 lineages with Lineages 1 and 2 being the most similar while Lineage 3 was clearly divergent (**Figure 5E**). Perhaps not surprising, THY ASCs from both female and male mice possessed the greatest percentage of Lineage 3 “only” cells as Lineage 3 possessed a large proportion of Cluster 0 (**Figure 5F**). For all lineages, proliferation genes such as *Mki67* (**Figure 5G**) showed an overall decrease over pseudotime which would be expected for ASCs transitioning from a cycling plasmablast state to a more mature, non-proliferative plasma cell existence. Lineages 1 and 2 demonstrated increased expression of key ASC regulatory genes such as *Jchain* (Hendrickson et al., 1995), *Mzb1* (Andreani et al., 2018; Rosenbaum et al., 2014) and *Cd74* (Pelletier et al., 2010) by the termination of pseudotime (**Figure 5G**). In contrast, Lineage 3 showed no such increase (**Figure 5G**). This pattern of gene expression was pervasive amongst the lineages (**Figures S5A-S5C**) and may be indicative of the ultimate fate of Lineages 1 and 2 being that of a mature, long-lived PC rather than Lineage 3 which may be destined for a shortened lifespan. In agreement with this idea, Lineage 3 also displayed downregulation of *Birc5* (**Figure S5C**) which encodes for Survivin, a key regulator of multiple myeloma growth and survival (Abdi et al., 2019; Romagnoli et al., 2007). Of unknown significance, the gene most highly associated with Lineage 3 and that demonstrated increased expression over pseudotime was the long non-coding RNA (lncRNA), *Gm42418* (**Figure S5C**). This lncRNA has been shown to associate with the NLRP3 inflammasome in macrophages following stimulation with lipopolysaccharide and nigericin (Zhang et al., 2019).

The above data showed that organ site could influence lineage composition within the total ASC pool. Thus, we were curious as to whether tissue localization would influence gene expression within the ASC clusters that we previously identified. To evaluate this, we examined gene expression changes within clusters as determined by organ (e.g., THY versus SPL) using a Log_2_ Fold Change > |0.5| and an Adjusted p-value < 0.05 as cut-offs for significance (**Tables S3-S5**). When comparing THY clusters to those from the SPL (**Figure 6A and Table S3**) and BM (**Figure 6B and Table S4**), majority of gene changes were present as downregulated in the THY. This was primarily limited to Clusters 0-3 which made up the bulk of THY ASCs. Female THY ASC clusters, in general, had increased numbers of differentially expressed genes when compared to males (**Figures 6A-6B and Tables S3-S4**). This trend was also present when differentially expressed genes between BM and SPL ASC clusters were examined (**Figure 6C and Table S5**).

**Figure 6.**
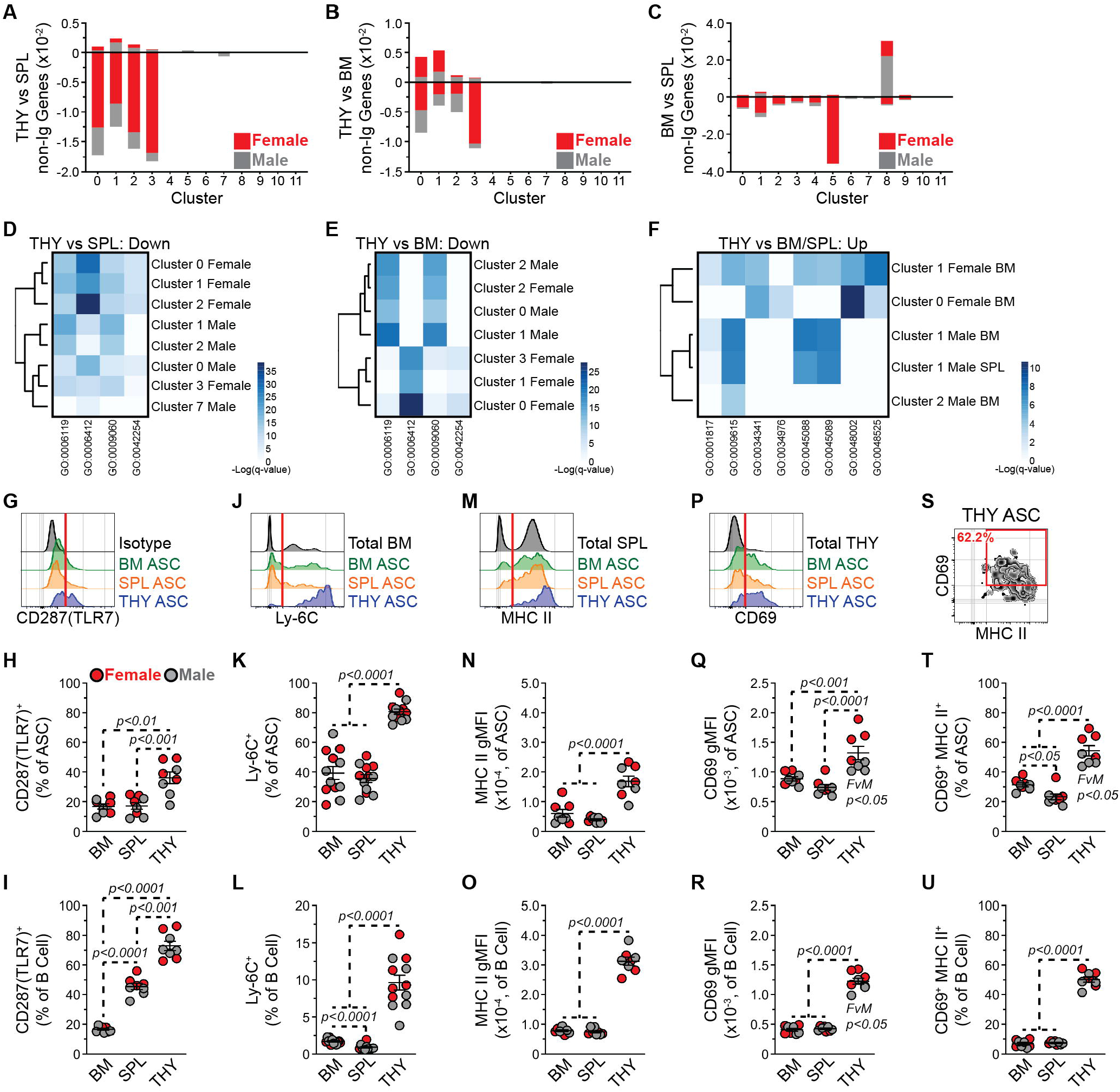
Thymus antibody-secreting cells possess transcriptional and protein indicators of immunological activation. (A-B) Number of non-Ig genes whose expression was increased (positive number) or decreased (negative number) in ASCs from the THY relative to those from (A) SPL and (B) BM. Data shown for females and males for all clusters. (C) Number non-Ig of genes whose expression was increased (positive number) or decreased (negative number) in ASCs from the BM relative to those from SPL. Data shown for females and males for all clusters. (D-F) Heatmaps depicting -Log(q-value) statistical significance for selected GO categories associated with THY clusters whose gene expression was (D) decreased relative to SPL, (E) decreased relative to BM and (F) increased relative to BM and/or SPL. Values derived from Metascape analyses presented in Tables S6-S7. (G) Representative flow cytometry histograms depicting intracellular CD287(TLR7) staining of BM, SPL and THY ASCs. Isotype control staining of BM ASCs is shown for comparison. Red vertical line added to CD287(TLR7) histograms to show cut-off for positive staining. (H-I) Percentages of CD287(TLR7)^+^ cells within (H) ASCs and (I) B220^+^ CD19^+^ B cells from BM, SPL and THY. (J) Representative flow cytometry histograms depicting Ly-6C surface staining of BM, SPL and THY ASCs. Total BM is shown for comparison. Red vertical line added to Ly-6C histograms to show cut-off for positive staining. (K-L) Percentages of Ly-6C^+^ cells within (K) ASCs and (L) B220^+^ CD19^+^ B cells from BM, SPL and THY. (M) Representative flow cytometry histograms depicting MHC II surface staining of BM, SPL and THY ASCs. Total SPL is shown for comparison. Red vertical line added to MHC II histograms to show cut-off for positive staining. (N-O) MHC II geometric mean fluorescence intensities (gMFIs) of (N) ASCs and (O) B220^+^ CD19^+^ B cells from BM, SPL and THY. (P) Representative flow cytometry histograms depicting CD69 surface staining of BM, SPL and THY ASCs. Total THY is shown for comparison. Red vertical line added to CD69 histograms to show cut-off for positive staining. (Q-R) CD69 gMFIs of (Q) ASCs and (R) B220^+^ CD19^+^ B cells from BM, SPL and THY. (S) Representative flow cytometry plot depicting CD69 and MHC II dual surface staining on THY ASCs. (T-U) Percentages of CD69^+^ MHC II^+^ cells within (T) ASCs and (U) B220^+^ CD19^+^ B cells from BM, SPL and THY. (H-I, K-L, N-O, Q-R, T-U) Symbols represent individual mice. Female: n = 4; Male: n = 4. Horizontal lines represent mean ± SEM. Unpaired Student’s t-Test for comparisons within sex. One-way ANOVA with Tukey’s correction for comparisons between organs.

Since we were particularly curious as to the difference between THY ASCs versus their SPL and BM counterparts, we performed gene ontology analyses (**Tables S6-S7**) similar to what was described above. In this instance, we decreased the threshold to 5 for the number of non-Ig genes included in a category. Genes downregulated in THY ASCs versus those of the SPL (**Figure 6D and Table S6**) and BM (**Figure 6E and Table S7**) were significantly enriched for categories involved in both protein production (e.g., GO:0006412, GO:42254) and metabolism (e.g., GO:0006119, GO:0009060). While there was some variation between clusters and sexes, these observations were generally applicable and may indicate THY ASCs being less “fit”. While the number of genes upregulated in clusters from THY ASCs was more limited, there was significant association with categories such as those related to antigen processing and presentation of peptide antigen (GO:0048002), response to interferon (IFN)-gamma (γ) (GO:0034341), regulation of innate immune response (GO:0045088) and response to virus (GO:0009615) among others (**Figure 6F and Tables S6-S7**). Most of these categories showed overlap of genes which included *Tlr7, Irf7* and a variety of IFN-stimulated genes (e.g., *Isg15*).

To confirm some of these findings, we performed intracellular flow cytometry (ICFC) on BM, SPL and THY ASCs for CD287(TLR7). This target was chosen in part due to its role in suppressing the reactivation of endogenous retroviruses which can lead to the development of T cell acute lymphoblastic leukemia (T-ALL) (Yu et al., 2012). CD287(TLR7) was detectable to some degree in ASCs from all 3 organs (**Figure 6G**). However, THY ASCs demonstrated a significantly higher percentage of CD287(TLR7)^+^ cells when compared to those from the BM and SPL (**Figure 6H**) with no significant differences observed between sexes. Total THY B cells (B220^+^ CD19^+^) also displayed an increased percentage of CD287(TLR7)^+^ cells (**Figure 6I**). Unlike with ASCs, SPL B cells possessed an intermediate phenotype relative to BM and THY (**Figure 6I**). Our gene ontology analyses suggested that THY ASC clusters possessed, at least to some degree, an IFN-inducible, or related, gene signature which is not all too surprising since CD287(TLR7) stimulation can induce IFN production (Saitoh et al., 2017). Along these lines, we consistently observed increased expression of *Ly6c2* in THY ASC clusters when compared to both SPL and BM clusters (**Tables S3-S4**). The *Ly6c2* gene product is part of the Ly-6C cell surface complex which can be induced by both Type I and Type II interferons (Jutila et al., 1988; Schlueter et al., 2001). In agreement with our scRNA-seq data, Ly-6C expression was significantly increased on the surface of THY ASCs (**Figures 6J-6K**). In fact, ∼81% of THY ASCs were Ly-6C^+^ compared to ∼39% and 36% of BM and SPL ASCs, respectively (**Figure 6K**). In contrast, only ∼10% of THY B cells expressed Ly-6C which was still shown to be an increase over those in the BM and SPL (**Figure 6L**). IFNγ signaling can also induce expression of major histocompatibility complex class II (MHC II) (Collins et al., 1984) and at least in females, we observed increased gene expression of *H2-Ab1* and *H2-Eb1* from THY Clusters 0 and 1 when compared to those from the BM. Using an Ab that recognizes I-A/I-E (MHC II), we performed flow cytometry and observed significantly increased MHC II surface expression on ASCs from THY as compared to those from BM and SPL (**Figures 6M-6N**). This was not unique to ASCs as THY B cells demonstrated a similar phenotype (**Figure 6O**) as previously reported (Perera et al., 2013). While THY ASCs demonstrated potential signs of IFN responsiveness, they also presented with evidence of overall cellular activation. In this case, CD69 demonstrated increased expression on the surface of THY ASCs (**Figures 6P-6Q**). In agreement with earlier studies (Ferrero et al., 1999), THY B cells were also hyper-activated as determined by CD69 gMFI (**Figure 6R**). Further evaluation of CD69 and MHC II showed a pattern of co-expression on the surface of THY ASCs (**Figure 6S**). Compared to those from BM and SPL, both THY ASCs (**Figure 6T**) and B cells (**Figure 6U**) demonstrated increased percentages of CD69^+^ MHC II^+^ cells. Taken together, these data revealed that our scRNA-seq dataset could be leveraged to identify THY ASC phenotypes related to various cell signaling/stimulatory events.

### Antibody-secreting cells display sex- and organ-specific differences in VDJ repertoire

Previous examination of differentially expressed genes based upon cluster, organ and sex all suggested Ig variance which was particularly evident at the *Ighv* gene family level. To better characterize these differences, we performed clonotype analysis on 10x Genomics VDJ-enriched libraries generated from BM, SPL and THY ASCs of female and male mice. High confidence contigs were generated and clonotypes were identified based upon Ig gene usage and complementarity-determining region (CDR) 3 nucleotide sequences using scRepertoire (Borcherding et al., 2020). Clonotypes were merged with the scGEX data and overlaid onto cluster UMAP plots (**Figure 7A**) which were originally derived in **Figure 5C**. BM and SPL ASCs possessed a bias towards representation from the left side of the UMAP which contained Clusters 2-6 and 9-11 (**Figure 7A**). In contrast, THY ASCs had the best overall representation across their total clusters (**Figure 7A**). When we examined clonality at the cluster level, it was obvious that clusters were differentially represented in our VDJ dataset (**Figure 7B**). For example, Clusters 3 and 9-10 demonstrated >50% detection of high confidence VDJ contigs within their respective ASC pools. In contrast, Clusters 0 and 7-8 demonstrated the least representation. Many of the clusters showed evidence of clonal expansion (i.e., hyperexpanded clones (0.1 < x ≤ 1)) which was not strictly an artifact of depth of clonal detection/assignment as detection of hyperexpanded clones was not dependent on percentages of ASCs with VDJ contigs per cluster (**Figure 7B**). We next assessed clonal similarity between clusters using the Jaccard similarity index (**Figure 7C**) in which higher numbers represented higher similarity. Overall, clusters appeared to be clonally distinct from one another. This was even true for Clusters 3 and 9-10 which were located in similar UMAP space (**Figure 5C**) and had similar depth of analysis within their respective ASC populations (**Figure 7B**). In this instance, Cluster 3 had similarity indices to Clusters 9 and 10 of 0.059 and 0.042, respectively (**Figure 7C**), while Clusters 9 and 10 had a similarity index of 0.018. For comparison, Clusters 5 and 6 demonstrated the highest similarity with a score of 0.287 (**Figure 7C**) on a scale from 0 to 1.

**Figure 7.**
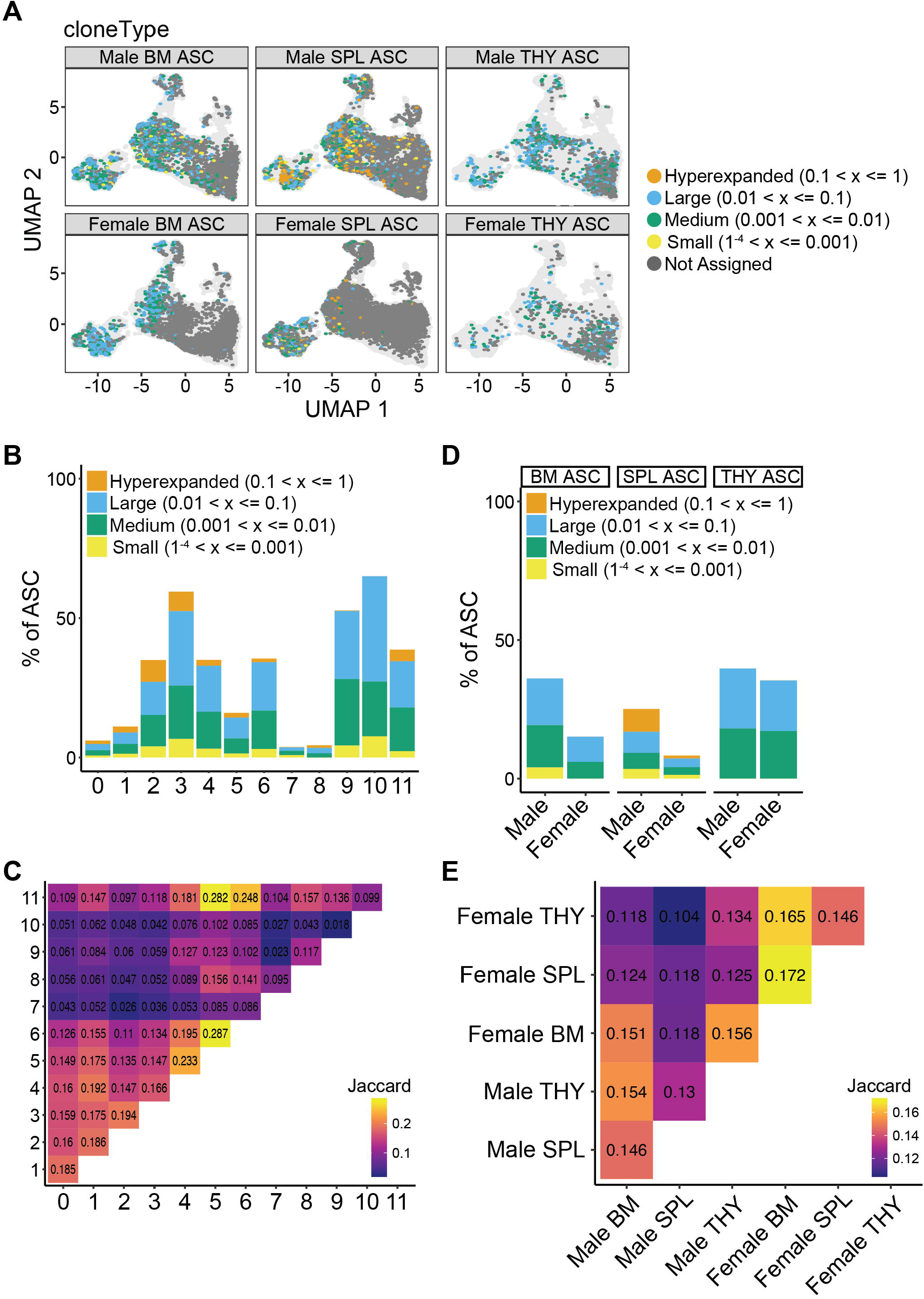
Antibody-secreting cell immunoglobulin repertories can be distinguished by both sex and organ. (A) Individual ASCs with high confidence Ig contigs overlaid upon cluster UMAPs from Figure 5C. Colors correspond to clone size. (B) Percentages of ASCs per cluster with high confidence Ig contigs as well as distribution of clone sizes within individual clusters. (C) Plot of Jaccard indices comparing similarities between repertoires from each cluster. (D) Percentages of ASCs per sex and organ with high confidence Ig contigs as well as distribution of clone sizes within each sample. (E) Plot of Jaccard indices comparing similarities between repertoires from each sex and organ.

Finally, we examined clonality based upon sample (i.e., organ and sex) (**Figures 7D-7E**). Variability regarding clonal sizes was observed when female and male ASCs were compared in either the BM or SPL. However, this may in part be due to the much higher clonotype representation in the male BM and SPL versus the female versions (**Figure 7D**). Within the THY, ASC clone sizes were mostly medium (0.001 < x ≤ 0.01) to large (0.01 < x ≤ 0.1) and consistent across sexes (**Figure 7D**). Similar to what was observed for clusters, there was minimal overlap between samples (**Figure 7E**). This was observed when organs from a particular sex were evaluated. For example, the male THY demonstrated similarity scores of 0.154 and 0.130 when compared to male BM and SPL, respectively. Additionally, similarity was low between sexes. This was on display for all 3 organs as female versus male BM, SPL and THY had Jaccard indices of 0.151, 0.118 and 0.134 (**Figure 7E**). These data suggested that the clonal repertoire of ASCs was dictated by the tissue of residence as well as by the sex in which those cells developed.

## Discussion

In the present study, we have provided a comprehensive resource which compares female and male ASCs found in the THY to their counterparts in organs such as the BM and SPL. By performing these comparisons, we have been able to identify characteristics that set THY ASCs apart and may lend themselves to unique cellular functions and regulatory mechanisms.

Upon initial assessment of ASC production, we observed significantly higher ASC numbers in young female mice relative to males. However, this sex-bias was limited to the THY and disappeared by middle-age as a result of the selective ASC expansion in the male THY combined with ASCs in the female THY having reached a carrying capacity by 3 mo. of age. This was in stark contrast to the BM and SPL, which demonstrated age-associated expansion of ASCs in both sexes albeit greater in female BM and SPL. This may simply be reflective of how ASCs in these different organs are generated over the course of aging. Through the use of intravenous Ab labeling and by blocking CD154(CD40L) signals in young mice, we demonstrated that THY ASCs were most likely generated locally and were dependent on T cell-derived signals for their generation. However, recent examination of LAG-3^+^ ASCs in the BM, SPL and mesenteric lymph node (mLN) demonstrated that at least some ASC populations gain independence from T cells for their generation over the course of aging (Dang et al., 2022). So why would this transition not take place in the THY? The answer may sit with what type of B cells reside in these organs as they age. Age-associated B cells (ABCs, CD19^+^ CD93^-^ CD43^-^ CD21^-^ CD23^-^) are typically thought of as being highly responsive to innate stimuli and have been shown to accumulate in both the SPL (Hao et al., 2011) and THY (Cepeda et al., 2018) of aging mice. However, distinct differences exist between these populations as ABCs in the SPL have been shown to retain IgM expression (Hao et al., 2011) while those in the THY are class-switched (Cepeda et al., 2018). This may be a predictor of different activation/signaling requirements which would agree with our *in vivo* CD40 agonism studies as well as previous *in vitro* studies comparing activation stimuli of THY and SPL B cells (Inaba et al., 1990; Inaba et al., 1995).

ASCs continue to gain prominence as an immunomodulatory population due in part to their ability to secrete immunosuppressive as well as inflammatory factors (Fillatreau, 2019; Pioli, 2019; Wang et al., 2021). Additionally, this same secretory ability may also influence their potential to regulate BM hematopoiesis (Meng et al., 2019; Pioli et al., 2019). As a result, understanding ASC biology has become of growing importance as the repertoire of ASC functions seems to depend on the context in which ASCs exist. Along these lines, recent studies have utilized RNA-seq at both bulk (Lino et al., 2018; Pioli et al., 2019; Price et al., 2019; Wilmore et al., 2021) and single cell (scRNA-seq) (Lam et al., 2018; Neumeier et al., 2022) levels in an attempt to define different ASC subsets or functional states. In general, these datasets have provided valuable resources that have led to the determination of critical ASC regulatory mechanisms. In this report, we compared ASCs from the female and male BM, SPL and THY using scRNA-seq. In some ways, our data were confirmatory to previous studies (Lam et al., 2018) which provided validity to our analyses. However, our results provided unique insights into THY ASCs and demonstrated their divergence from their BM and SPL counterparts. Via analysis of developmental trajectory (i.e., pseudotime), we defined 3 ASC lineages with the 3^rd^ of these appearing to be unfit and potentially destined for a shortened lifespan as exemplified by reduced expression of genes related to ASC function (e.g., *Jchain, Mzb1, Cd74*) and survival (e.g., *Birc5*). Our data in conjunction with previous studies seemed to indicate the derivation of THY ASCs from potentially autoreactive THY B cells (Perera et al., 2013; Perera et al., 2016; Yamano et al., 2015). As such, it is preferential that THY ASCs were enriched for Lineage 3 as this may be an evolutionary failsafe to prevent the long-standing presence of autoreactive ASCs. In this regard, it would be interesting to transcriptionally profile THY resident autoreactive ASCs that exist in diseases such as myasthenia gravis (Hill et al., 2008). Would their gene signature resemble that of Lineages 1 and 2 rather than Lineage 3?

Regarding a potential function(s) for THY ASCs, our transcriptional profiling revealed significant anti-viral and interferon gene ontology enrichment within genes specifically upregulated in THY ASC clusters. This included genes that code for protein products including TLR7, Ly-6C and MHC II, the latter being a key factor in antigen presentation. Accordingly, our ASC phenotyping demonstrated immunoglobulin staining on the surface of THY ASC. As such, it is tempting to speculate that autoreactive ASCs could internalize self-antigen present in the THY microenvironment and present that same self-antigen to selecting T cells, thus playing a role previously attributed to THY B cells (Perera et al., 2013; Perera et al., 2016; Yamano et al., 2015). There is precedent for this behavior as ASCs in the LN have been shown to present antigen to T follicular helper cells and suppress their behavior following model antigen immunization (Pelletier et al., 2010). Additionally, this would potentially explain our observations regarding the minimal overlap of immunoglobulin clonotypes in THY ASCs compared to those in the BM and SPL. However, comparison of THY T cell receptor (TCR) repertoires and autoreactivity in ASC wildtype and knockout mice would be needed to further support a role for ASCs in THY negative selection. Alternatively, the same TCR comparisons performed in animals in which ASCs lack components of the antigen presentation machinery (e.g., MHC II) would provide a pathway specific analysis.

Finally, aside from the initial skewing of THY ASC numbers, we did observe additional sex-biases. These manifested mostly within the immunoglobulin repertoire but could also be observed when ASC phenotypes were examined. For example, THY ASCs from female mice expressed higher levels of CD69 on the cell surface and possessed increased percentages of CD69^+^ MHC II^+^ cells within their total ASC pool. While the significance of this remains to be determined, these observations coincide with increased overall THY output in young female versus male mice (Gui et al., 2011; Min et al., 2006) and overall increased immunoreactivity that has been observed in the female sex (Klein and Flanagan, 2016).

In summary, the data presented here support the concept that THY ASCs develop independently of their peripheral counterparts and possess a gene signature that is dictated by their local microenvironment (e.g., interferon signaling in the THY (Yu et al., 2012)). Importantly, we have developed a comprehensive scRNA-seq data resource that can be utilized by researchers to develop testable hypotheses investigating the development, maintenance and function of THY ASCs.

## Supporting information

Key Resources

## Acknowledgments

Funding was provided by the Western Michigan University Homer Stryker M.D. School of Medicine (WMed) via intramural startup funds. Flow cytometry was performed at the WMed Flow Cytometry and Imaging Core. The authors thank WMed students Michael Crone, Matthew Kornas and Patrick Renner for their technical assistance. The authors thank the Van Andel Genomics Core and Bioinformatics and Biostatistics Core for providing sequencing facilities and analysis services, respectively.

## Author Contributions

K.T.P and P.D.P designed experiments. K.T.P. and P.D.P. conducted and analyzed experiments. K.H.L. performed bioinformatics analyses. P.D.P wrote the manuscript and all authors approved of the manuscript.

## STAR Methods

### Contact for Reagent and Resource Sharing

Further information and requests for resources and reagents should be directed to and will be fulfilled by the Lead Contact, Peter Dion Pioli.

## EXPERIMENTAL MODEL AND SUBJECT DETAILS

### Experimental Animals

Mouse strains B6.Cg-Tg(Prdm1-EYFP)1Mnz/J (Stock #: 008828), B6.129-*Prdm1*^tm1Clme/^J (Stock #: 008100) and B6.C(Cg)-*Cd79a*^tm1(cre)Reth/^EhobJ (Stock #: 020505) were purchased from the Jackson Laboratory. Mice were considered young at 3-4 mo. of age. Mice were considered middle-aged at 12 mo. of age. Female and male mice were utilized for all experiments. Animal care and use were conducted according to the guidelines of the Western Michigan University Homer Stryker M.D. School of Medicine (WMed) Institutional Animal Care and Use Committee (IACUC). All animals were housed and/or bred in the WMed vivarium.

## METHOD DETAILS

### Isolation of Bone Marrow, Spleen and Thymus Tissue

All tissues were processed and collected in calcium and magnesium-free 1x phosphate buffered saline (PBS). Spleen (SPL) and thymus (THY) were dissected and crushed between the frosted ends of two slides. Bone marrow (BM) was isolated from both femurs and tibias by cutting off the end of bones and flushing the marrow from the shaft using a 23-gauge needle. Cell suspensions were centrifuged for 5 minutes at 4 °C and 600g. Red blood cells were lysed by suspending cells in 3 mL of 1x red blood cell lysis buffer on ice for ∼3 minutes. Lysis was stopped with the addition of 7 mL of 1x PBS. Cell suspensions were counted in a hemocytometer using Trypan Blue to exclude dead cells and passed through 40-μm filters before use.

Whole blood was obtained via cardiac puncture and immediately transferred to EDTA containing tubes on ice. 100 μL was directly mixed with 3 mL of 1x red blood cell lysis buffer and placed on ice for ∼3 minutes. Lysis was stopped with the addition of 7 mL of 1x PBS.

### Immunostaining

All staining procedures were performed in 1x PBS + 0.1% bovine serum albumin (BSA). Samples were labeled with a CD16/32 blocking antibody (Ab) to eliminate non-specific binding of Abs to cells via Fc receptors. All Abs utilized are listed in the Key Resources Table. For surface staining, cells were incubated on ice for 30 minutes with the appropriate Abs. Unbound Abs were washed from cells with 1x PBS + 0.1% BSA followed by centrifugation for 5 minutes at 4 °C and 600g. Supernatants were decanted and cell pellets were resuspended in an appropriate volume of 1x PBS + 0.4% BSA + 2 mM EDTA for flow cytometric analysis. Before analysis, cells were strained through a 40 μM filter mesh.

Intracellular CD287(TLR7) staining was performed using the Foxp3/Transcription Factor Staining Buffer Set. Cells were surface stained as described above. Afterwards, cell pellets were resuspended in 1 mL of buffer #1 and incubated at room temperature (RT) for 30 minutes in the dark. Subsequently, 2 mL of buffer #2 were added and cells centrifuged for 5 minutes at RT and 600g. Supernatants were discarded and cells were blocked with total mouse IgG at RT in the dark for 20-30 minutes. Cells were then incubated with either anti-CD287(TLR7)-PE or mouse IgG1-PE isotype control Abs. Additional samples were left unstained for comparison. Following this incubation, 2 mL of buffer #2 were added and cells centrifuged for 5 minutes at RT and 600g. Finally, cells were washed with 3 mL of 1x PBS + 0.1% BSA and centrifuged for 5 minutes at RT and 600g. Cells were prepped for flow cytometric analysis as described above. For all intracellular stains, eBioscience Fixable Viability (Live-Dead) Dye eFluor 780 (Thermo Fisher Scientific, Catalog # 65-0865-14) was added to samples to assess dead cell content. The stock solution was diluted 1:250 and 10 μL was added to ∼5 × 10^6^ cells per stain. Live-Dead stain was added concurrent with surface staining Abs.

### Flow Cytometry and Fluorescence-Activated Cell Sorting

Flow cytometry was performed on a Fortessa (BD Biosciences) located in the Flow Cytometry and Imaging Core at WMed. Data were analyzed using FlowJo (v10) software. Viable cells were gated using side scatter area (SSC-A) versus forward scatter (FSC-A) area. Singlets were identified using sequential gating of FSC-width (W) versus FSC-A, SSC-W versus SSC-A and FSC-height (H) versus FSC-A.

Fluorescence-activated cell sorting (FACS) was performed on a Melody (BD Biosciences) located in the Flow Cytometry and Imaging Core at WMed. Prior to sorting, cells were resuspended at a final concentration of 2×10^7^ cells per mL in αMEM supplemented with HEPES (25 mM), Penicillin-Streptomycin (100 U/mL), L-glutamine (2 mM), Gentamicin (50 μg/mL) and EDTA (2 mM) and subsequently passed through a 40 μM filter mesh. Viable cells were gated using SSC-A versus FSC-A. Singlets were identified using sequential gating of FSC-W versus FSC-A, SSC-W versus SSC-A and FSC-H versus FSC-A.

### Enzyme-Linked Immunosorbent Spot Assay

Enzyme-linked immunosorbent spot (ELISpot) plates (Millipore Sigma, Cat# MSIPS4W10) were briefly incubated at RT for 1 minute with 15 μL of 35% ethanol (EtOH). EtOH was removed and wells were washed 3 times with 150 μL of 1x PBS. Subsequently, wells were coated overnight (O/N) at 4 °C with 100 μL of capture Ab. The capture Ab (Millipore Sigma, Cat# SAB3701043-2MG) recognized mouse total IgG+IgM+IgA isotypes and was pre-diluted in 1x PBS to a final concentration of 5 μg/mL before use. The next day, coating Ab was removed and wells were washed 3 times with 150 μL RPMI 1640. Plates were subsequently blocked with 150 μL RPMI for a minimum of 2 hours at 37 °C in a 5% CO_2_/20% O_2_ tissue culture incubator. Blocking solution was removed and 100 μL of RPMI supplemented with a proliferation-inducing ligand (APRIL) (10 ng/mL), interleukin (IL)-6 (10 ng/mL), heat-inactivated fetal calf serum (10%), Penicillin-Streptomycin (100 U/mL), L-glutamine (2 mM), Gentamicin (50 μg/mL), sodium pyruvate (1 mM), non-essential amino acids (1x), non-essential vitamins (1x) and 2-mercaptoethanol (10^−05^ M). Antibody-secreting cells (ASCs) from THY were directly FACS-purified into wells with a target number of 100 cells per well (in triplicate). Cells were then incubated O/N (>12 hours) at 37 °C in a 5% CO_2_/20% O_2_ tissue culture incubator. The next day, culture supernatants and cells were removed. Wells were washed 3 times with 150 μL of 1x PBS then an additional 3 times with 150 μL of 1x PBS + 0.1% Tween-20 + 1% BSA. Secondary Abs conjugated to horse radish peroxidase (HRP) were added at a volume of 100 μL per well and plates were incubated for 2 hours at RT. Anti-IgM-HRP (Thermo Fisher Scientific, Cat# 62-6820) and anti-IgG-HRP (SouthernBiotech, Cat# 1015-05) were diluted 1:25,000 and 1:50,000 in 1x PBS + 0.1% Tween-20 + 1% BSA before use, respectively. Following incubation, Abs were removed plates washed 3 times with 150 μL of 1x PBS + 0.1% Tween-20 + 1% BSA. An additional 3 washes with 150 μL of 1x PBS were performed. To reveal “spots”, 100 μL of Developing Solution from the AEC Substrate Set (BD Biosciences, Cat# 551951) was added to each well. Plates were shaken at 200 rpm for 30 minutes at RT. Developing Solution was removed and plates were washed 5 times with 150 μL of H_2_O. Well backing was removed and plates were dried at RT after which “spots” were visualized with a CTL ELISPOT reader. Using Adobe Photoshop, auto-contrast was uniformly applied to wells and brightness was set to 100 for visualization.

### *In Vivo* CD45 Labeling

Young (3-4 mo. old) mice received either 1x PBS or αCD45-PE (1 μg) antibody in a 100 μL volume via intravenous (i.v.) injection into the retro-orbital (r.o.) sinus. Mice were euthanized 5 minutes or 24 hours post-injection for analysis.

### *In Vivo* CD154(CD40L) Blockade

Young (3-4 mo. old) mice received a total of 1200 μg of hamster IgG isotype control or anti-mouse CD154(CD40L) (MR-1) antibodies. Administration was split evenly amongst 12 total intraperitoneal (i.p.) injections (100 μg per injection, 100 μl volume) over a 4-week span after which mice were euthanized for analysis. Antibodies were supplied by Bio X Cell and pre-diluted to 1000 μg/mL using InVivoPure pH 7.0 Dilution Buffer.

### *In Vivo* CD40 Stimulation

On days -1 and 0, young (3-4 mo. old) mice were administered 100 μg (100 μl volume) of rat IgG2a isotype control or agonistic anti-mouse CD40 antibodies per injection via the i.p. route. Antibodies were supplied by Bio X Cell and pre-diluted to 1000 μg/mL using InVivoPure pH 7.0 Dilution Buffer. Mice were euthanized for analysis on days 2 and 28 following the last injection.

### 10x Genomics Single Cell Library Preparation, Sequencing and Analysis

ASCs from BM, SPL and THY were FACS-purified and collected into 1x PBS + 0.1% BSA. Recovery was verified by hemocytometer count and/or analysis via flow cytometry. Cells were concentrated in a volume of 40 μL following centrifugation for 10 minutes at 4 °C and 600g. A volume of 37.8 μL of cells was loaded into a Chromium Next GEM Chip G and single cell gel emulsions were generated on a 10x Genomics Chromium Controller. 10x Genomics barcoded full-length cDNA was then synthesized on a Veriti 96-Well Thermal Cycler (Thermo Fisher Scientific) following the standard 10x Genomics protocol. Following initial recovery and cleanup, cDNA was further amplified 18 cycles based upon manufacturer recommendations.

V(D)J-enriched libraries and gene expression (GEX) libraries were generated using the Chromium Single Cell V(D)J Enrichment Kit, Mouse B Cell (10x Genomics, Cat# 1000072) and Chromium Next GEM Single Cell 5’ Library and Gel Bead Kit v1.1 (10x Genomics, Cat# 1000165), respectively. All libraries were sample indexed for multiplexing with the Chromium i7 Multiplex Kit (10x Genomics, Cat# 120262) following standard 10x Genomics protocols. All completed libraries were visualized and quantified using an Agilent 2100 Bioanalyzer. Furthermore, qPCR utilizing the Collibri Library Quantification Kit (Thermo Fisher Scientific, Cat# A38524500) was performed to verify the suitability of each library for sequencing. V(D)J libraries were pooled in tandem based upon organ and population (e.g., Female and Male THY ASC) and sequenced on an Illumina MiSeq inhouse. All 6 GEX libraries were pooled and sequenced simultaneously on a NovaSeq 6000 at the Van Andel Institute Genomics Core.

Reads were aligned to the mm10 reference genome (‘refdata-gex-mm10-2020-A’ provided by 10x) to obtain UMI counts using the cellranger count pipeline v6.0.2 with default parameters. The filtered counts matrices, with non-cell-associated barcodes removed, were imported into R v4.1.0 as a SingleCellExperiment object using DropletUtils v1.12.2 (Lun et al., 2019). Low total UMI counts or number of genes detected indicate low-quality cells while a high percent of reads mapping to mitochondrial genes suggest lysed cells (Ilicic et al., 2016). To filter by these metrics, median absolute deviations (MAD) for these metrics were calculated for each sample separately and 3 MAD from the median was used as a cutoff, as implemented in the ‘perCellQCMetrics’ and ‘quickPerCellQC’ functions in the scuttle package v1.2.0 (Amezquita et al., 2020; McCarthy et al., 2017). A total of 3122 cells, with 1359 from the female bone marrow sample, were removed due to high mitochondrial gene expression but none were removed based on the other two metrics.

The filtered SingleCellExperiment object was converted to a Seurat (v4.0.3) object for ‘integration’ (batch correction) across the samples and for cluster identification using the standard Seurat integration workflow (Stuart et al., 2019). Briefly, the Seurat object was split based on the sample. UMI counts were normalized for total expression in each cell and log-transformed. The top 2000 most variable genes within each sample were identified, from which the ‘anchor’ genes for integration were chosen using the ‘SelectIntegrationFeatures’ function. The ‘FindIntegrationAnchors’ and ‘IntegrateData’ functions were used to perform batch correction using a method that utilizes mutual nearest neighbors (MNN) to identify biologically correspondent cells between datasets (Stuart et al., 2019). The corrected expression values were scaled and used for Principal Component Analysis (PCA). The top 30 PCs were chosen for downstream analyses based on an elbow plot. For visualization, Uniform Manifold Approximation and Projection (UMAP) was performed on the top 30 PCs. Similarly, a shared nearest neighbor graph of the cells was constructed using the top 30 PCs and used for unsupervised clustering using the Louvain algorithm. Clustering was done at multiple resolutions ranging from 0.1 to 0.9, but the 0.3 resolution clustering was chosen for subsequent analyses. Various UMAP plots and bar charts were plotted using the dittoSeq v1.4.1 package (Bunis et al., 2020).

Marker genes for clusters were identified using the ‘findMarkers’ function in scran v1.20.1 with the parameters, ‘pval.type=“any”‘, ‘direction=“up”‘, ‘test.type=“wilcox”‘, ‘min.prop=0.75’ and setting block to the sample. This performs pairwise comparisons of each cluster to every other cluster, testing for up-regulation using Wilcoxon rank sum tests in each sample (block) separately; the p-values for each gene across all block levels are combined using Stouffer’s Z-score method and the reported AUC is a weighted average of the AUCs across all the block levels. For each gene and cluster, the summary effect size is the effect size from the pairwise comparison with the lowest p-value and the combined p-value is computed using Simes’ method. The ‘min.prop’ parameter affects only the ‘Top’ column in the results. With ‘min.prop’ set to 0.75, genes with Top <= N are top N marker genes in at least 0.75 of the comparisons. Differentially expressed genes between samples within each cluster were identified using the ‘FindMarkers’ function in Seurat with ‘logfc.threshold’ set to 0.5. Only comparisons where both groups had greater than 20 cells were tested.

Pseudotime trajectories were inferred on the UMAP embedding using the ‘slingshot’ function in the Slingshot v2.0.0 package (Street et al., 2018), with the parameter ‘approx_points=150’ and setting the root at cluster 11, which had high expression of proliferative marker genes. To test for genes associated with pseudotime, the tradeSeq v1.6.0 package (Van den Berge et al., 2020) was used. For genes with a count of at least 3 in at least 100 cells, a negative binomial GAM was fitted using the ‘fitGAM’ function, providing the batches (sample) using the ‘U’ argument. Pseudotime-associated genes were identified using the ‘associationTest’ function. Smoothed expression along pseudotime was plotted for various genes using the functions ‘predictSmooth’ and ‘plotSmoothers’.

For VDJ analysis, reads were aligned to the mm10 VDJ reference provided by 10x (‘refdata-cellranger-vdj-GRCm38-alts-ensembl-5.0.0’) and assembled into contigs using the cellranger vdj pipeline v6.0.2 with default parameters. The ‘filtered_contig_annotations.csv’ files were read into R and combined for use with the scRepertoire v1.3.5 R package (Borcherding et al., 2020) using the ‘combineBCR’ function. Clonotype information was merged with the Seurat object using the ‘combineExpression’ function; this also calculated clonotype frequencies within each sample. Overlap of clonotypes was assessed by the Jaccard index using the ‘clonalOverlap’ function with the parameters, ‘cloneCall=“gene+nt”‘ and ‘method=“jaccard”‘.

Code is available at: https://github.com/vari-bbc/Pioli_2022_ASCs_code. Sequencing data has been deposited in the NCBI Gene Expression Omnibus (GEO) database and is accessible under accession number GSE193701.

## QUANTIFICATION AND STATISTICAL ANALYSIS

The numbers of mice used or replicates performed per experiment are listed in the figure legends. For non-genomics data, statistical analyses were performed using GraphPad Prism (v8.4.2) software. A Student’s t-Test was used for statistical comparisons between 2 groups. For comparisons among 3 or more groups, a one-way ANOVA with Tukey’s or Dunnett’s correction was utilized. Both paired and unpaired tests were utilized where appropriate. Statistically significant p-values are shown within each figure.

## Supplemental Information Titles and Legends

**Figure S1. Thymus antibody-secreting cells express expected surface markers. Related to Figure 1**.

(A) Total BM cell numbers. (B) Percentages of BM PBs and PCs. (C) Numbers of BM PBs and PCs. (D) Total SPL cell numbers. (E) Percentages of SPL PBs and PCs. (F) Numbers of SPL PBs and PCs. (A-F) Symbols represent individual mice. Female: n = 11; Male: n = 11. Horizontal lines represent mean ± SEM. Unpaired Student’s t-Test for inter-sex comparisons. Paired Student’s t-Test for intra-sex comparisons. (G-K) Flow cytometry geometric mean fluorescence intensities (gMFIs) of (G) CD267(TACI), (H) CD44, (I) Sca-1(LY-6A/E), CD184(CXCR4) and mIgκ+λ on the surface of PBs and PCs from BM, SPL and THY. Symbols represent individual mice. Horizontal lines represent mean ± SEM. One-way ANOVA with Tukey’s correction. (G-H, K) Female: n = 3; Male: n = 4. (I-J) Female: n = 6; Male: n = 6.

**Figure S2. Fluorescence-activated cell sorting of antibody-secreting cells. Related to Figures 1 and 5**.

(A) Representative flow cytometry plots depicting live cell and singlet gating based upon various SSC and FSC parameters. (B) Representative flow cytometry plots illustrating gating of ASCs from BM, SPL and THY. ASCs were defined as CD138^HI^ IgD^-/LO^ CD90.2^-/LO^ Prdm1-eYFP^+^.

**Figure S3. Quantification of antibody-secreting cell numbers in aging bone marrow, spleen and thymus. Related to Figure 1**.

(A) Middle-aged (12 mo. old) female and male Prdm1-eYFP mice were assayed for ASCs BM, SPL and THY. Middle-aged data compared to that of young (3 mo.) mice generated in Figure 1. (B) Total THY cell numbers. (C-D) Percentages of THY (B) PBs and (D) PCs. (E-F) Numbers of THY (E) PBs and (F) PCs. (G) Total BM cell numbers. (H-I) Percentages of BM (H) PBs and (I) PCs. (J-K) Numbers of BM (J) PBs and (K) PCs. (L) Total SPL cell numbers. (M-N) Percentages of SPL (M) PBs and (N) PCs. (O-P) Numbers of SPL (O) PBs and (P) PCs. (A-P) Symbols represent individual mice. Female 3 mo.: n = 11; Male 3 mo.: n = 11; Female 12 mo.: n = 6; Male 12 mo.: n = 9. Horizontal lines represent mean ± SEM. Unpaired Student’s t-Test.

**Figure S4. Gene ontology analysis of antibody-secreting cell clusters. Related to Figure 5**.

(A) Numbers of cluster marker genes that were Ig-related or non-Ig-related. Ig-related genes consisted of those encoding either V, D and J segments or constant regions for heavy and light chains. (B-E) Heatmaps depicting -Log(q-value) statistical significance for cluster marker gene association with selected (B) Protein Production, (C) Cellular Metabolism, (D) RNA and (E) Cell Cycle & Survival GO categories. Values derived from Metascape analyses presented in Table S2. (F) UMAP plot showing log-normalized unique molecular identifier (UMI) counts for *Ccna2*.

**Figure S5. Antibody-secreting cell pseudotime lineages demonstrate progressive changes in gene expression. Related to Figure 5**.

(A-C) Heatmaps showing row-normalized, smoothed gene expression of the top 100 lineage-associated genes for (A) Lineage 1, (B) Lineage 2 and (C) Lineage 3. Pseudotime progresses from left-to-right.

**Table S1. Marker genes for each cluster. Related to Figure 5**.

**Table S2. Metascape gene ontology analysis of non-Ig cluster marker genes. Related to Figure 5.**

**Table S3. Differentially expressed genes between thymus and spleen clusters. Related to Figure 6**.

**Table S4. Differentially expressed genes between thymus and bone marrow clusters. Related to Figure 6**.

**Table S5. Differentially expressed genes between bone marrow and spleen clusters. Related to Figure 6**.

**Table S6. Metascape gene ontology analysis of non-Ig cluster genes differentially expressed between thymus and spleen. Related to Figure 6**.

**Table S7. Metascape gene ontology analysis of non-Ig cluster genes differentially expressed between thymus and bone marrow. Related to Figure 6**.

