## Supplementary material for "Thymus Antibody-Secreting Cells Represent a Tissue Specific Population That Possess an Activated Cellular Phenotype": Figure S1

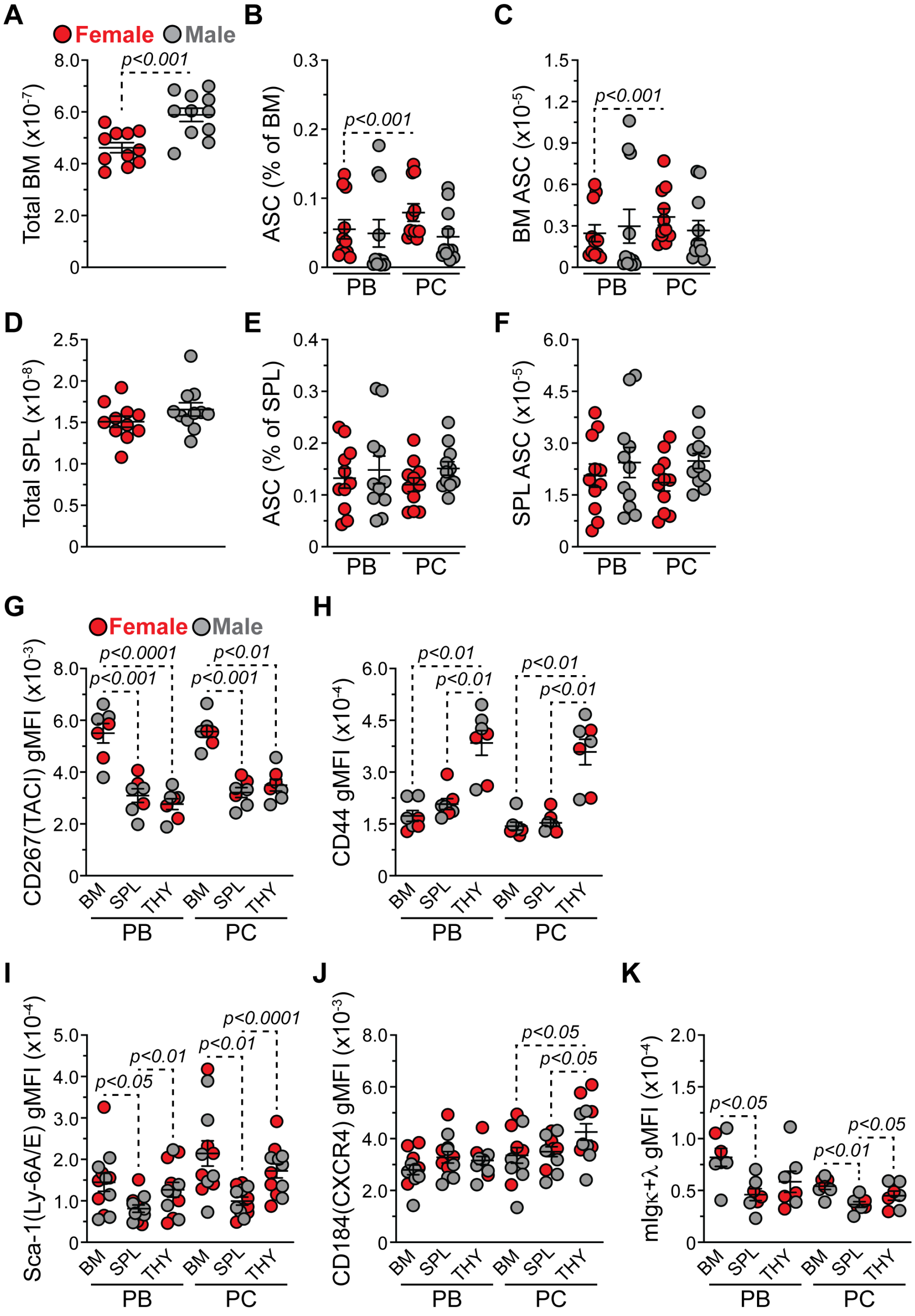


**Figure S1. Thymus antibody-secreting cells express expected surface markers. Related to Figure 1.**

(A) Total BM cell numbers. (B) Percentages of BM PBs and PCs. (C) Numbers of BM PBs and PCs. (D) Total SPL cell numbers. (E) Percentages of SPL PBs and PCs. (F) Numbers of SPL PBs and PCs. (A-F) Symbols represent individual mice. Female: n = 11; Male: n = 11. Horizontal lines represent mean ± SEM. Unpaired Student's t-Test for inter-sex comparisons. Paired Student's t-Test for intra-sex comparisons. (G-K) Flow cytometry geometric mean fluorescence intensities (gMFIs) of (G) CD267(TACI), (H) CD44, (I) Sca-1(LY-6A/E), CD184(CXCR4) and mIgκ+λ on the surface of PBs and PCs from BM, SPL and THY. Symbols represent individual mice. Horizontal lines represent mean ± SEM. One-way ANOVA with Tukey's correction. (G-H, K) Female: n = 3; Male: n = 4. (I-J) Female: n = 6; Male: n = 6.
