## Supplementary material for "Thymus Antibody-Secreting Cells Represent a Tissue Specific Population That Possess an Activated Cellular Phenotype": Figure S2

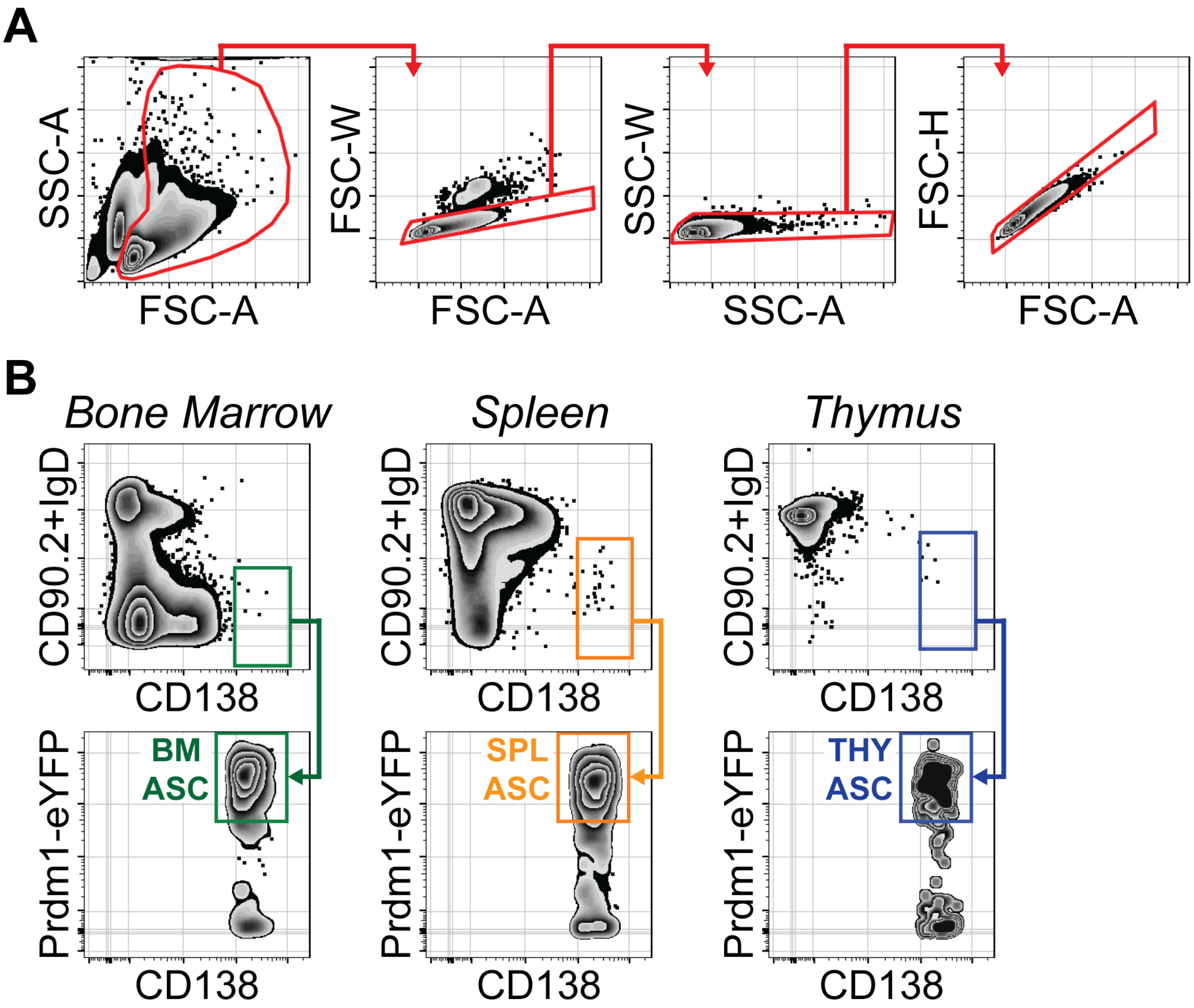


**Figure S2. Fluorescence-activated cell sorting of antibody-secreting cells. Related to Figures 1 and 5.**

(A) Representative flow cytometry plots depicting live cell and singlet gating based upon various SSC and FSC parameters. (B) Representative flow cytometry plots illustrating gating of ASCs from BM, SPL and THY. ASCs were defined as $CD138^{HI}$ $IgD^{-/LO}$ $CD90.2^{-/LO}$ Prdm1-$eYFP^{+}$.
