## Supplementary material for "Thymus Antibody-Secreting Cells Represent a Tissue Specific Population That Possess an Activated Cellular Phenotype": Figure S3

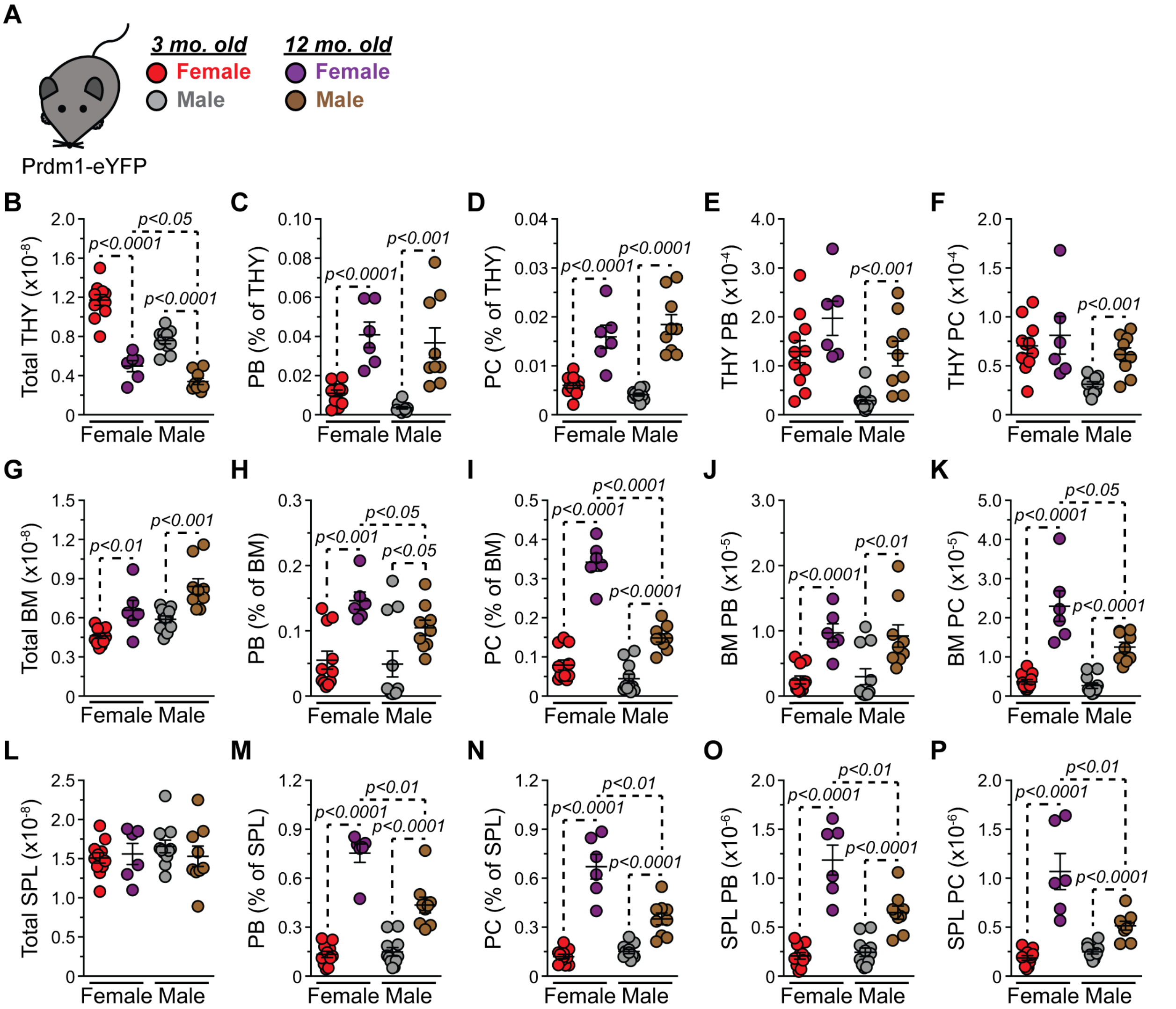


**Figure S3. Quantification of antibody-secreting cell numbers in aging bone marrow, spleen and thymus. Related to Figure 1.**

(A) Middle-aged (12 mo. old) female and male Prdm1-eYFP mice were assayed for ASCs BM, SPL and THY. Middle-aged data compared to that of young (3 mo.) mice generated in Figure 1. (B) Total THY cell numbers. (C-D) Percentages of THY (B) PBs and (D) PCs. (E-F) Numbers of THY (E) PBs and (F) PCs. (G) Total BM cell numbers. (H-I) Percentages of BM (H) PBs and (I) PCs. (J-K) Numbers of BM (J) PBs and (K) PCs. (L) Total SPL cell numbers. (M-N) Percentages of SPL (M) PBs and (N) PCs. (O-P) Numbers of SPL (O) PBs and (P) PCs. (A-P) Symbols represent individual mice. Female 3 mo.: n = 11; Male 3 mo.: n = 11; Female 12 mo.: n = 6; Male 12 mo.: n = 9. Horizontal lines represent mean ± SEM. Unpaired Student's t-Test. Comparisons between young female and mice populations omitted for clarity.
