## Supplementary material for "Thymus Antibody-Secreting Cells Represent a Tissue Specific Population That Possess an Activated Cellular Phenotype": Figure S4

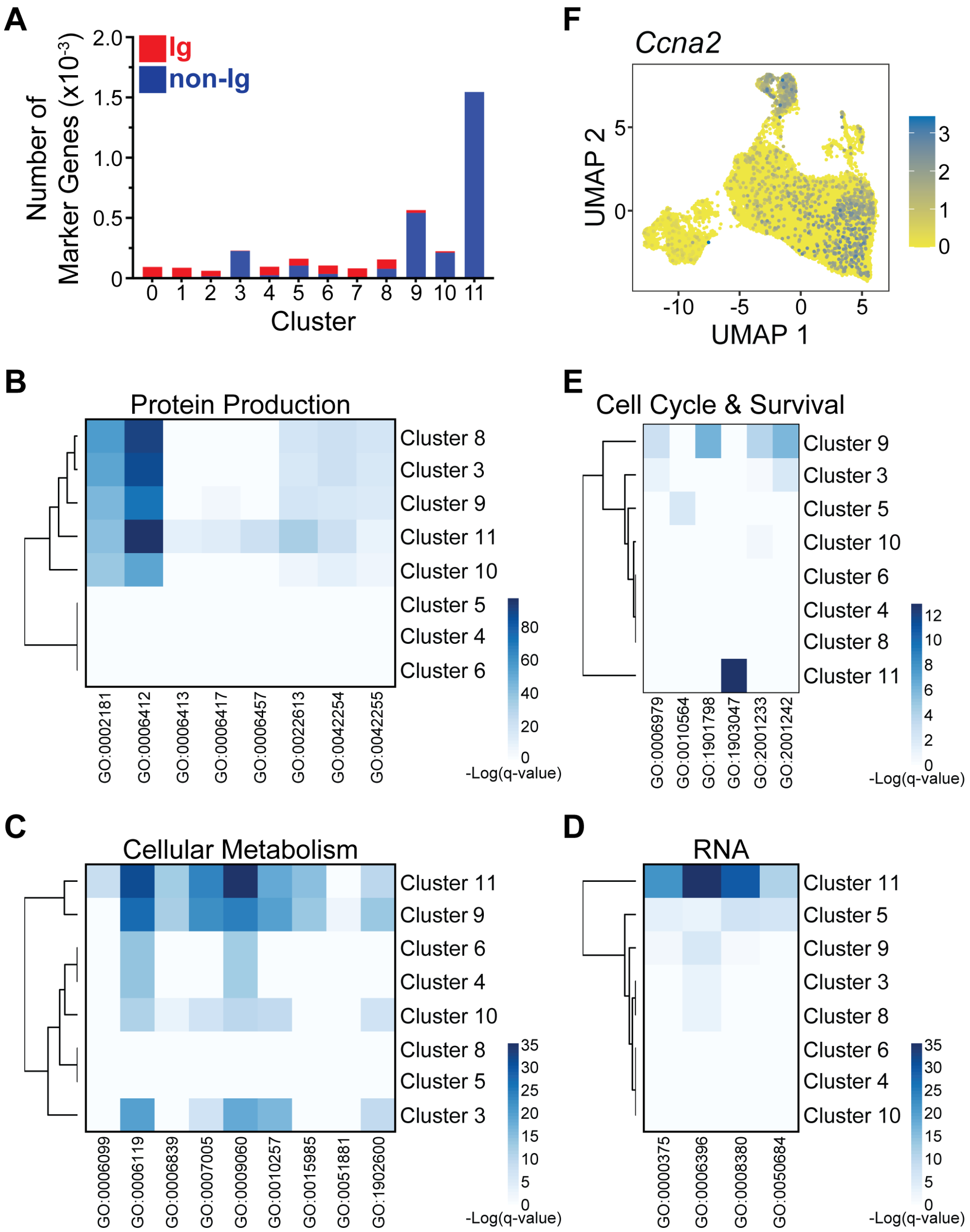


**Figure S4. Gene ontology analysis of antibody-secreting cell clusters. Related to Figure 5.** (A) Numbers of cluster marker genes that were Ig-related or non-Ig-related. Ig-related genes consisted of those encoding either V, D and J segments or constant regions for heavy and light chains. (B-E) Heatmaps depicting -Log(q-value) statistical significance for cluster marker gene association with selected (B) Protein Production, (C) Cellular Metabolism, (D) RNA and (E) Cell Cycle & Survival GO categories. Values derived from Metascape analyses presented in Table S2. (F) UMAP plot showing log-normalized unique molecular identifier (UMI) counts for *Ccna2*.
