## Supplementary material for "Thymus Antibody-Secreting Cells Represent a Tissue Specific Population That Possess an Activated Cellular Phenotype": Figure S5

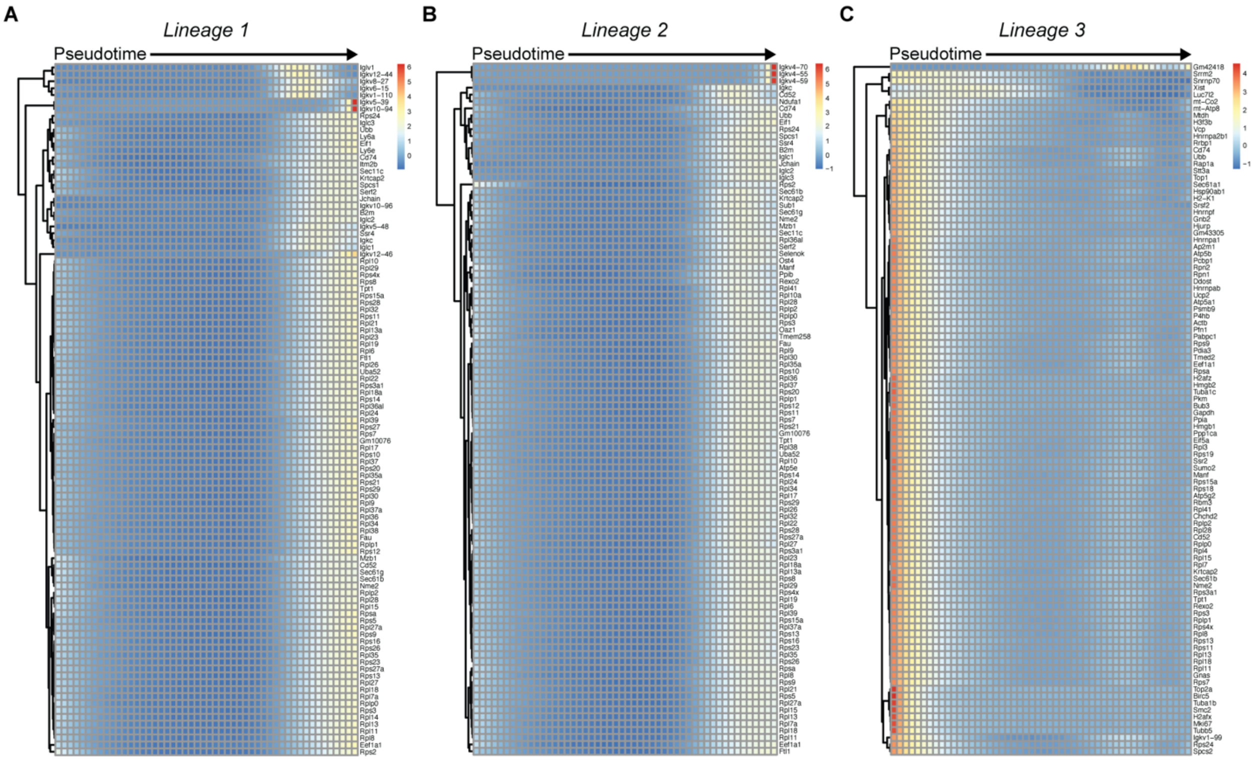


**Figure S5. Antibody-secreting cell pseudotime lineages demonstrate progressive changes in gene expression. Related to Figure 5.**

(A-C) Heatmaps showing row-normalized, smoothed gene expression of the top 100 lineage-associated genes for (A) Lineage 1, (B) Lineage 2 and (C) Lineage 3. Pseudotime progresses from left-to-right.
