## Supplementary material for "Thymus Antibody-Secreting Cells Represent a Tissue Specific Population That Possess an Activated Cellular Phenotype": Key Resources

**KEY RESOURCES TABLE**

The table highlights the genetically modified organisms and strains, cell lines, reagents, software, and source data **essential** to reproduce results presented in the manuscript. Depending on the nature of the study, this may include standard laboratory materials (i.e., food chow for metabolism studies), but the Table is **not** meant to be comprehensive list of all materials and resources used (e.g., essential chemicals such as SDS, sucrose, or standard culture media don’t need to be listed in the Table). **Items in the Table must also be reported in the Method Details section within the context of their use.** The number of **primers and RNA sequences** that may be listed in the Table is restricted to no more than ten each. If there are more than ten primers or RNA sequences to report, please provide this information as a supplementary document and reference this file (e.g., See Table S1 for XX) in the Key Resources Table.

***Please note that ALL references cited in the Key Resources Table must be included in the References list.*** Please report the information as follows:

- **REAGENT or RESOURCE:** Provide full descriptive name of the item so that it can be identified and linked with its description in the manuscript (e.g., provide version number for software, host source for antibody, strain name). In the Experimental Models section, please include all models used in the paper and describe each line/strain as: model organism: name used for strain/line in paper: genotype. (i.e., Mouse: OXTR^fl/fl^: B6.129(SJL)-Oxtr^tm1.1Wsy/J^). In the Biological Samples section, please list all samples obtained from commercial sources or biological repositories. Please note that software mentioned in the Methods Details or Data and Software Availability section needs to be also included in the table. See the sample Table at the end of this document for examples of how to report reagents.
- **SOURCE:** Report the company, manufacturer, or individual that provided the item or where the item can obtained (e.g., stock center or repository). For materials distributed by Addgene, please cite the article describing the plasmid and include “Addgene” as part of the identifier. If an item is from another lab, please include the name of the principal investigator and a citation if it has been previously published. If the material is being reported for the first time in the current paper, please indicate as “this paper.” For software, please provide the company name if it is commercially available or cite the paper in which it has been initially described.
- **IDENTIFIER:** Include catalog numbers (entered in the column as “Cat#” followed by the number, e.g., Cat#3879S). Where available, please include unique entities such as [RRIDs](https://www.force11.org/group/resource-identification-initiative), Model Organism Database numbers, accession numbers, and PDB or CAS IDs. For antibodies, if applicable and available, please also include the lot number or clone identity. For software or data resources, please include the URL where the resource can be downloaded. Please ensure accuracy of the identifiers, as they are essential for generation of hyperlinks to external sources when available. Please see the Elsevier [list of Data Repositories](https://www.elsevier.com/authors/author-resources/research-data/data-base-linking) with automated bidirectional linking for details. When listing more than one identifier for the same item, use semicolons to separate them (e.g. Cat#3879S; RRID: AB_2255011). If an identifier is not available, please enter “N/A” in the column.
  - ***A NOTE ABOUT RRIDs:*** We highly recommend using RRIDs as the identifier (in particular for antibodies and organisms, but also for software tools and databases). For more details on how to obtain or generate an RRID for existing or newly generated resources, please [visit the RII](https://www.force11.org/group/resource-identification-initiative) or [search for RRIDs](https://scicrunch.org/resources).

Please use the empty table that follows to organize the information in the sections defined by the subheading, skipping sections not relevant to your study. Please do not add subheadings. To add a row, place the cursor at the end of the row above where you would like to add the row, just outside the right border of the table. Then press the ENTER key to add the row. Please delete empty rows. Each entry must be on a separate row; do not list multiple items in a single table cell. Please see the sample table at the end of this document for examples of how reagents should be cited.

***TABLE FOR AUTHOR TO COMPLETE***

*Please upload the completed table as a separate document.* ***Please do not add subheadings to the Key Resources Table.*** *If you wish to make an entry that does not fall into one of the subheadings below, please contact your handling editor. (****NOTE:*** *For authors publishing in Current Biology, please note that references within the KRT should be in numbered style, rather than Harvard.)*

**KEY RESOURCES TABLE**

| REAGENT or RESOURCE | SOURCE | IDENTIFIER |
| --- | --- | --- |
| Antibodies | | |
| CD138-BV421 (Clone: 281-2) | BD Biosciences | Cat# 562610; RRID: AB_11153126 |
| IgD-BV605 (Clone: 11-26c.2a) | BioLegend | Cat# 405727; RRID: AB_2562887 |
| IgD-APC (Clone: 11-6c.2a) | BioLegend | Cat# 405714; RRID: AB_10643423 |
| CD90.2(Thy-1.2)-BV605 (Clone: 53-2.1) | BD Biosciences | Cat# 563008; RRID: AB_2665477 |
| CD90.2(Thy-1.2)-APC (Clone: 53-2.1) | BD Biosciences | Cat# 553007; RRID: AB_398526 |
| CD45R(B220)-PerCP/Cy5.5 (Clone: RA3-6B2) | BD Biosciences | Cat# 552771; RRID: AB_394457 |
| CD45R(B220)-BV711 (Clone: RA3-6B2) | BD Biosciences | Cat# 563892; RRID: AB_2738470 |
| CD19-BUV395 (Clone: 1D3) | BD Biosciences | Cat# 563557; RRID: AB_2722495 |
| CD16/32-Unlabeled (Clone: 93) | Thermo Fisher Scientific | Cat# 14-0161-86; RRID: AB_467135 |
| CD44-PE/Cy7 (Clone: IM7) | Thermo Fisher Scientific | Cat# 25-0441-81; RRID: AB_469622 |
| CD267(TACI)-PE (Clone: 8F10) | BioLegend | Cat# 133403; RRID: AB_2203542 |
| CD69-PE/Cy7 (Clone: H1.2F3) | BD Biosciences | Cat# 561930; RRID: AB_10893591 |
| I-A/I-E(MHC II)-AF647 (Clone: M5/114.15.2) | BioLegend | Cat# 107618; RRID: AB_493525 |
| Ly-6C-PE (Clone: HK1.4) | BioLegend | Cat# 128008; RRID: AB_1186132 |
| CD287(TLR7)-PE (Clone: A04B10) | BD Biosciences | Cat# 565557; RRID: AB_2739295 |
| Mouse IgG1-PE isotype control (Clone: MOPC-21) | BD Biosciences | Cat# 554680; RRID: AB_395506 |
| CD45-PE (Clone: 30F11) | BioLegend | Cat# 103106; RRID: AB_312971 |
| CD45-APC (Clone: 30F11) | BioLegend | Cat# 103112; RRID: AB_312977 |
| Igκ-PE (Clone: RMK-45) | BioLegend | Cat# 409505; RRID: AB_2563580 |
| Igλ-PE (Clone: RML-42) | BioLegend | Cat# 407308; RRID: AB_1027659 |
| Ly-6A/E(Sca-1)-PE/Cy7 (Clone: D7) | BioLegend | Cat# 108113; RRID: AB_493597 |
| CD184(CXCR4)-APC (Clone: L276F12) | BioLegend | Cat# 146507; RRID: AB_2562784 |
| Mouse IgG-Unlabeled | SouthernBiotech | Cat# 0107-01; RRID: AB_2732898 |
| Anti-mouse IgG+IgA+IgM (H+L) | Millipore Sigma | Cat# SAB3701043-2MG |
| Goat anti-mouse IgM-HRP | Thermo Fisher Scientific | Cat# 62-6820; RRID: AB_2533954 |
| Goat anti-mouse IgG-HRP | SouthernBiotech | Cat# 1015-05; RRID: AB_2794194 |
| InVivoMab polyclonal Armenian hamster IgG isotype control | Bio X Cell | Cat# BE0091; RRID: AB_1107773 |
| InVivoMab anti-mouse CD154(CD40L) (Clone: MR-1) | Bio X Cell | Cat# BE0017-1; RRID: AB_1107601 |
| InVivoMab rat IgG2a isotype control (Clone: 2A3) | Bio X Cell | Cat# BE0089; RRID: AB_1107769 |
| InVivoMab anti-mouse CD40 (Clone: FGK4.5/FGK45) | Bio X Cell | Cat# BE0016-2; RRID: AB_1107647 |
| Chemicals, Peptides, and Recombinant Proteins | | |
| InVivoPure pH 7.0 Dilution Buffer | Bio X Cell | Cat# IP0070 |
| Bovine Serum Albumin (DNase- and Protease-free) | Fisher Scientific | Cat# BP9706100 |
| APRIL | Fisher Scientific | Cat# 7907-AP-010CF |
| IL-6 | Fisher Scientific | Cat# PMC0066 |
| Critical Commercial Assays | | |
| Platinum II Host-Start PCR Master Mix (2x) | Thermo Fisher Scientific | Cat# 14000012 |
| Chromium Next GEM Single Cell 5' Library and Gel Bead Kit v1.1 | 10x Genomics | Cat# 1000165 |
| Chromium Next GEM Chip G Single Cell Kit | 10x Genomics | Cat# 1000120 |
| Chromium Single Cell V(D)J Enrichment Kit, Mouse B Cell | 10x Genomics | Cat# 1000072 |
| Chromium i7 Multiplex Kit | 10x Genomics | Cat# 120262 |
| Beckman Coulter SPRIselect | Fisher Scientific | Cat# NC0406406 |
| Dynabeads MyOne Silane | Thermo Fisher Scientific | Cat# 37002D |
| Collibri Library Quantification Kit | Thermo Fisher Scientific | Cat# A38524500 |
| High Sensitivity DNA Kit | Agilent | Cat# 5067-4626 |
| AEC Substrate Set | BD Biosciences | Cat# 551951; RRID: AB_2868954 |
| eBioscience Fixable Viability (Live-Dead) Dye eFluor 780 | Thermo Fisher Scientific | Cat# 65-0865-14 |
| Foxp3/Transcription Factor Staining Buffer Set | Thermo Fisher Scientific | Cat# 00-5523-00 |
| Deposited Data | | |
| ASC scRNA-Seq GEX and VDJ data | This paper | GEO: GSE193701 |
| Experimental Models: Organisms/Strains | | |
| Mouse: B6.Cg-Tg(Prdm1-EYFP)1Mnz/J | The Jackson Laboratory | JAX: 008828; RRID: IMSR_JAX:008828 |
| Mouse: B6.129-*Prdm1*^tm1Clme/^J | The Jackson Laboratory | JAX: 008100; RRID: IMSR_JAX:008100 |
| Mouse: B6.C(Cg)-*Cd79a*^tm1(cre)Reth/^EhobJ | The Jackson Laboratory | JAX: 020505; RRID: IMSR_JAX:020505 |
| Oligonucleotides | | |
| *Prdm1-eYFP* Genotyping Forward Primer: TTCCACAGCTCTGAGGGTCT | The Jackson Laboratory | https://www.jax.org/Protocol?stockNumber=008828&protocolID=25522 |
| *Prdm1-eYFP* Genotyping Reverse Primer: CGGTGGTGCAGATGAACTT | The Jackson Laboratory | https://www.jax.org/Protocol?stockNumber=008828&protocolID=25522 |
| *Prdm1* Floxed Genotyping Forward Primer:  CAATGCTTGTCTAGTGTC | The Jackson Laboratory | https://www.jax.org/Protocol?stockNumber=008100&protocolID=22773 |
| *Prdm1* Floxed Genotyping Reverse Primer:  AGTAGTTGAATGGGAGC | The Jackson Laboratory | https://www.jax.org/Protocol?stockNumber=008100&protocolID=22773 |
| *Mb1* (Cd79a) Genotyping Wildtype Forward Primer:  CTCTTTACCTTCCAAGCACTGA | The Jackson Laboratory | https://www.jax.org/Protocol?stockNumber=029412&protocolID=20206 |
| *Mb1* (Cd79a) Genotyping Common Reverse Primer:  ACTGAGGCAGGAGGATTGG | The Jackson Laboratory | https://www.jax.org/Protocol?stockNumber=029412&protocolID=20206 |
| *Mb1* (Cd79a) Genotyping Mutant Forward Primer:  CATTTTCGAGGGAGCTTCA | The Jackson Laboratory | https://www.jax.org/Protocol?stockNumber=029412&protocolID=20206 |
| Software and Algorithms | | |
| FlowJo (v10) | BD Biosciences | RRID: SCR_008520 |
| GraphPad Prism 8 | GraphPad Software | RRID: SCR_002798 |
| Adobe Illustrator | Adobe | RRID: SCR_010279 |
| Adobe Photoshop | Adobe | RRID: SCR_014199 |
| Cell Ranger v6.0 | 10x Genomics | https://support.10xgenomics.com/single-cell-gene-expression/software/downloads/latest |
| DropletUtils v1.12.2 | Lun et al., 2019 | https://bioconductor.org/packages/release/bioc/html/DropletUtils.html |
| scuttle v1.2.0 | Amezquita et al., 2020; McCarthy et al., 2017 | https://bioconductor.org/packages/release/bioc/html/scuttle.html |
| Seurat v4.0.3 | Stuart et al., 2019 | https://satijalab.org/seurat/ |
| dittoSeq v1.4.1 | Bunis et al., 2020 | https://bioconductor.org/packages/devel/bioc/vignettes/dittoSeq/inst/doc/dittoSeq.html |
| scran v1.20.1 | Lun et al., 2016 | https://bioconductor.org/packages/release/bioc/html/scran.html |
| Slingshot v2.0.0 | Street et al., 2018 | https://bioconductor.org/packages/release/bioc/html/slingshot.html |
| tradeSeq v1.6.0 | Van den Berge et al., 2020 | https://www.bioconductor.org/packages/release/bioc/html/tradeSeq.html |
| scRepertoire v1.3.5 | Borcherding et al., 2020 | http://www.bioconductor.org/packages/release/bioc/vignettes/scRepertoire/inst/doc/vignette.html |
